# Hypoxia-related immune subsets induced by *Salmonella* Typhi infection link early bacterial gut invasion to human infection outcomes

**DOI:** 10.1101/2025.01.08.631852

**Authors:** Noa Bossel Ben-Moshe, Shelly Hen Avivi, Liron Levy Efrati, Leia Veinman, Jennifer Hill, Daniel O’Connor, Marije Verheul, Lisa Stockdale, Florence McLean, Andrew Pollard, Roi Avraham

**Affiliations:** Department of Immunology and Regenerative Biology, Weizmann Institute of Science, Israel; Department of Paediatrics, Oxford Vaccine Group, University of Oxford, and the NIHR Oxford Biomedical Research Centre, UK

## Abstract

*Salmonella* Typhi (*S.* Typhi), the causative agent of typhoid disease, remains a major public health concern. Owing to the human-restricted nature of *S*. Typhi, current studies of typhoid pathogenesis in animal models are limited to a murine non-typhoidal pathogen. Furthermore, human studies are limited to analyses of peripheral immune responses which are blind to tissue-specific immunity and do not allow perturbations. What is now needed is an integrative approach that will provide mechanistic insights into *S*. Typhi pathogenesis and immune correlates of infection outcome. Here, we performed an integrated single-cell analysis of immune responses from the human *S.* Typhi challenge model and mouse model of typhoid disease, to associate biological mechanism with human infection outcome. Most prominent, we revealed immune subsets with a hypoxia-related signature in circulating immune cells from individuals that develop disease in the human challenge model. This signature was also evident in the mouse model in activated macrophages infiltrating into the Peyer’s patches, but not during infection with a mutant strain impaired for gut invasion. We further identified hypoxia-related signature as a general immune correlate of disease outcome in other infection- and inflammatory-related diseases. Collectively, using integrated analysis of mouse and human infection models, we revealed a hypoxia-related signature that link immune responses during bacterial invasion to increased risk of developing typhoid disease in humans, suggesting a possible causative role during the development of typhoid disease.

## Introduction

Typhoid fever is a major public health concern, with an estimated nine million cases and 110,000 deaths annually^1^. Typhoid disease is caused by ingestion of *Salmonella enterica* serovar Typhi (*S.* Typhi), a host-restricted pathogen whose reservoir is humans. Unlike other *Salmonella* serovars that primarily cause local intestinal inflammation and diarrhea, *S.* Typhi invades from the gastrointestinal tract into the bloodstream, survives and reproduces within macrophages, and can result in chronic carriage^2^. *S.* Typhi was declared by the WHO as one of the priority pathogens requiring the development of new treatments to stop the spread of antimicrobial resistance^3^. However, owing to the human-restricted nature of *S*. Typhi, current studies of typhoid pathogenesis in animal models are limited to a murine pathogen. Furthermore, human studies provide insight only to the peripheral protective immune responses which are blind to tissue-specific immunity and mechanistic insight via perturbations.

Our understanding of typhoid disease progression is mostly afforded by the mouse typhoid fever model, which uses a murine non-typhoidal strain. In this model, susceptible mice carrying a mutation in the *Nramp1* gene are infected orally with the closely related *Salmonella enterica* serotype Typhimurium (*S.* Typhimurium) without streptomycin pretreatment, resulting in a systemic infection that mimics typhoid disease^4,5^. Using this model, the route of bacterial dissemination from the gut to systemic sites has been closely monitored. Following oral infection, *S.* Typhimurium enters the small intestine, traverses the epithelial barrier through microfold cells (M cells), and invades Peyer’s patches (PP) located in the terminal ileum. The bacteria then disseminate to lymph nodes and the bloodstream, ultimately reaching systemic organs, similar to typhoid-like disease^6,7^. The mouse model of typhoid fever has been invaluable for identifying major bacterial virulence components required for systemic infection, such as *aroA*^8^ and *htrA*^9^, for elucidating interactions with the microbiota^10,11^, and for studying *Salmonella*-specific immune responses in the intestinal mucosa^10,12–14^. Importantly, similar routes of infection and immune responses in mice were also reported for other gut-invading pathogens, such as *Listeria monocytogenes* and *Yersinia pseudotuberculosis*^15–18^. However, evidence directly linking immune activation in the mouse model to the human immune responses to *S*. Typhi infection is still lacking. More importantly, the immune correlates of infection outcomes in humans cannot be studied in the mouse model.

To overcome these limitations, human *S.* Typhi challenge studies have been established^19–21^. In these studies, healthy volunteers are orally exposed to *S.* Typhi under controlled conditions and closely monitored. Participants either develop typhoid disease or remain asymptomatic, with all participants receiving antibiotics 14 days after bacterial ingestion or at the time of disease onset. Analyzing blood samples during the course of infection can facilitate the investigation of immune correlates of protection. For instance, multifunctional CD8+ T cells are linked to protective immune responses^22^, while CD8+ T memory cells^23^ and NK cell activity with IFNγ release^24^ are associated with increased disease susceptibility. During typhoid disease onset, CD8+ mucosal-associated invariant T (MAIT) cells decrease^25^, and intermediate monocytes increase^26^. Transcriptome profiling of whole-blood (WB) and peripheral blood mononuclear cells (PBMCs) from these studies has identified significant cytokine elevation^27^ and monocyte activation^28^ as early as 12 hours post-bacterial ingestion, although these early responses do not correlate with infection outcomes. Acute disease is characterized by marked elevation in Type I and II interferon (IFN) responses, which are associated with bacteremia^27^ and elevation of monocyte activation markers^26^. While these studies capture significant changes in peripheral blood immune responses during *S.* Typhi infection, they lack insight into the rich biology and variety of specialized immune cells at the site of infection.

Advances in single-cell RNA-seq (scRNA-seq) have enabled detailed investigation of immune responses at the level of individual cells within complex tissues and samples. It is now becoming increasingly understood that during infection, tissue-specific immune subsets drive local host-pathogen interactions that lead to different infection outcomes. For example, during early systemic infection, *S.* Typhimurium is found in the spleen within permissive CD9^+^ macrophages expressing an anti-inflammatory gene program, including fatty acid and lipid metabolism mediated by PPAR-γ^29^. During chronic infection, bacterial control in spleen granulomas is mediated by two distinct populations: ACE^+^ macrophages and iNOS^+^ macrophages which are more bacterial-permissive^30^. ScRNA-seq has also been instrumental in analyzing the immune responses of PBMCs in human challenge models. For example, early immune responses associated with protective immunity or disease progression were described in human challenge models of West Nile virus^31^, SARS-CoV2^32^, and influenza^33^. The challenge now is to bridge findings from animal models and human challenge models, enabling a unified view of the infection process from tissue-specific immune responses at the site of infection to systemic disease outcomes.

In this study, using scRNA-seq, we set out to study immune subsets from the human typhoid challenge model and from the gut tissue in a mouse model of typhoid disease. We profiled PBMCs from individuals in the *S.* Typhi human challenge study with different disease outcomes at different time points after the challenge. We observed a hypoxia-related gene signature in several immune subsets enriched specifically from individuals who developed the disease during the study, already several days before the onset of symptoms. To link our findings and gain mechanistic insight into tissue-specific immunity, we turned to survey the mouse typhoid model. In the gut tissue, we identified changes mainly in the PP as a key site for bacterial colonization. The main changes included infiltration of activated macrophages with a hypoxia-related signature, identical to the hypoxia-related signature in humans. To functionally analyze this hypoxia-related gene signature, we infected mice with an *S.* Typhimurium mutant strain impaired for PP invasion. We found that the hypoxia signature was dependent on the pathogen invasion and colonization of the PP. Finally, we found this hypoxia-related gene signature additionally correlated with sepsis and inflammatory bowel disease patient outcomes, hypoxic cutaneous leishmaniasis lesions in humans, and colonization of the PP with *Yersinia pseudotuberculosis* in mice. Thus, we identified an early hypoxic signature of immune cells that precedes the development of human typhoid disease and is linked to bacterial gut invasion.

## Results

### Transcriptional changes of blood immune cells in response to human oral infection with *S.* **Typhi**

To characterize human immune subset changes in response to *S.* Typhi infection, we obtained blood samples from individuals of the human challenge model^21^. In this cohort, healthy volunteers were orally infected with *S.* Typhi (Quailes strain) and monitored for 14 days, during which time blood samples were collected at pre-defined time points. To assess the transcriptional changes of blood immune cells in individuals with different disease outcomes, we selected six individuals from this study: three who developed typhoid disease (TD), as indicated by clinical symptoms of fever or positive blood culture, and three individuals who did not develop typhoid disease (nTD). All individuals received antibiotic treatment at the end of the study. We performed scRNA-seq on Peripheral Blood Mononuclear Cells (PBMCs) collected from these six individuals at three time points: prior to the challenge (T0), twelve hours post-challenge (T12), and at the time of disease diagnosis for those who developed typhoid disease (TD), or seven days post-challenge for those who remained healthy (T7d; the median time point for disease onset) (**Fig. 1a**). Overall, we sequenced 165,415 cells from all samples together. We clustered the cells and classified them according to cell type marker genes into natural killer (NK) cells (NKG7 and GNLY), T cells (CD3D/G), B cells (CD79A/B and MS4A1), monocytes and macrophages (LYZ and MS4A4), and dendritic cells (DCs; FSCN1) (**Fig. 1b, Supplementary Fig. 1a, and Supplementary Table 1**). Following *S.* Typhi challenge, we observed infiltration of monocytes into the bloodstream and a reduction in T cell numbers across all individuals (**Supplementary Fig. 1b**). The t-distributed Stochastic Neighbor Embedding (t-SNE) projection of the data into two-dimensional space illustrated changes in cell states following *S.* Typhi challenge (**Fig. 1c**), between individuals (**Fig. 1d**), and between disease outcomes (**Fig. 1e-f**). To quantitatively determine these observed variations in cell states between conditions, we calculated integrated Local Inverse Simpson’s Index (iLISI) scores to assess the mixing of cells across individuals and disease outcomes^34^. This metric measures the mixture level of each cell across its local neighbors on the tSNE graph (from **Fig. 1d-e**). Using this measurement, we were able to quantitively estimate the inter-individual compositions of immune cell states relevant to infection (**Fig. 1g**). We observed that while there are individual-specific immune states, others are common across multiple individuals (general response). Additionally, we observed outcome-specific immune states that are exclusive to individuals with the same clinical outcome. Thus, in addition to the expected inter-individual variability in the immune responses to *S.* Typhi infection, we observed outcome-specific immune responses.

**Figure 1:**
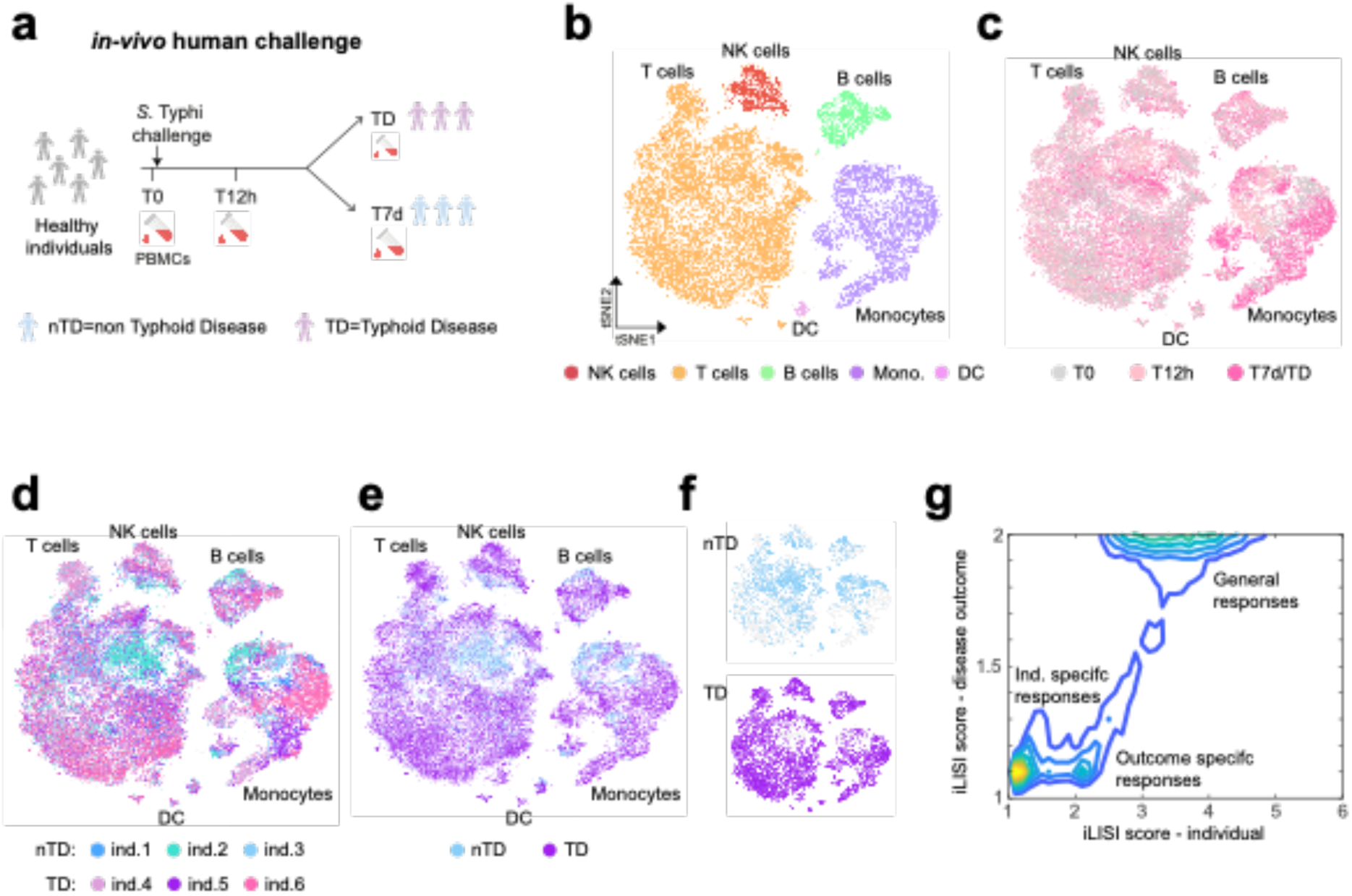
scRNA-seq analysis of PBMCs from individuals in the *S.* Typhi human challenge study. **(a)** Overview of the study design and cohort for the scRNA-seq experiment: PBMC samples were collected from six individuals in the *S.* Typhi human challenge study, including three who developed typhoid disease (TD) and three who did not (nTD). Samples were taken at three time points for each individual: before challenge (T0), twelve hours post-challenge (T12h), and at the time of disease diagnosis for TD individuals (TD) or seven days post-challenge for nTD individuals (T7d). **(b-f)** tSNE visualization of scRNA-seq data combining all individuals and time points together. Cells are colored by cell type in **(b)**, time point in **(c)**, individual in **(d)**, and disease outcome (nTD or TD) in **(e)**. **(f)** tSNE plot from (e), separated to show immune cells from nTD individuals (top) and TD individuals (bottom) alone, highlighting outcome-specific subsets. Cell-type annotations and color legends are provided. **(g)** Density plot of iLISI scores for each cell, measuring the degree of mixing across individuals (x-axis) and disease outcomes (y-axis). iLISI scores close to 1 indicate no mixing, while higher scores indicate greater mixing. Three types of immune cell responses are highlighted: 1) Individual-specific responses (iLISI scores close to 1 for both individuals and outcomes mixing), 2) General responses (iLISI scores > 3 for individuals and close to 2 for outcomes), and 3) Outcome-specific responses (iLISI score close to 1 for outcomes and > 2 for individuals).

### Monocyte and macrophage response dynamics correlate with disease outcome

The identification of immune states that are specific to disease outcomes motivated us to determine the specific subsets and states that differ between individuals who developed typhoid disease and those who did not. To this end, we focused on the responses of monocytes and macrophages, as these are the primary target cells for *S.* Typhi infection. First, we subclustered monocytes and macrophages to the subset level (**Fig. 2a**). Overall, we identified 15 monocyte and macrophage subsets. The marker genes for each subset included cell type markers, immune-related genes, and metabolic-related genes (**Supplementary Fig. 2a and Supplementary Table 2**), allowing us to functionally annotate these subsets. Notably, specific subsets were linked to infection outcomes (**Fig. 2b**). Among these, a CD9/CXCR4 subset, an M1 subset (pro-inflammatory state of macrophages), and a subset associated with glycolysis and hypoxia markers, were all linked with typhoid disease. Non-classical monocytes and type I IFN response were associated with individuals who did not develop the disease.

**Figure 2:**
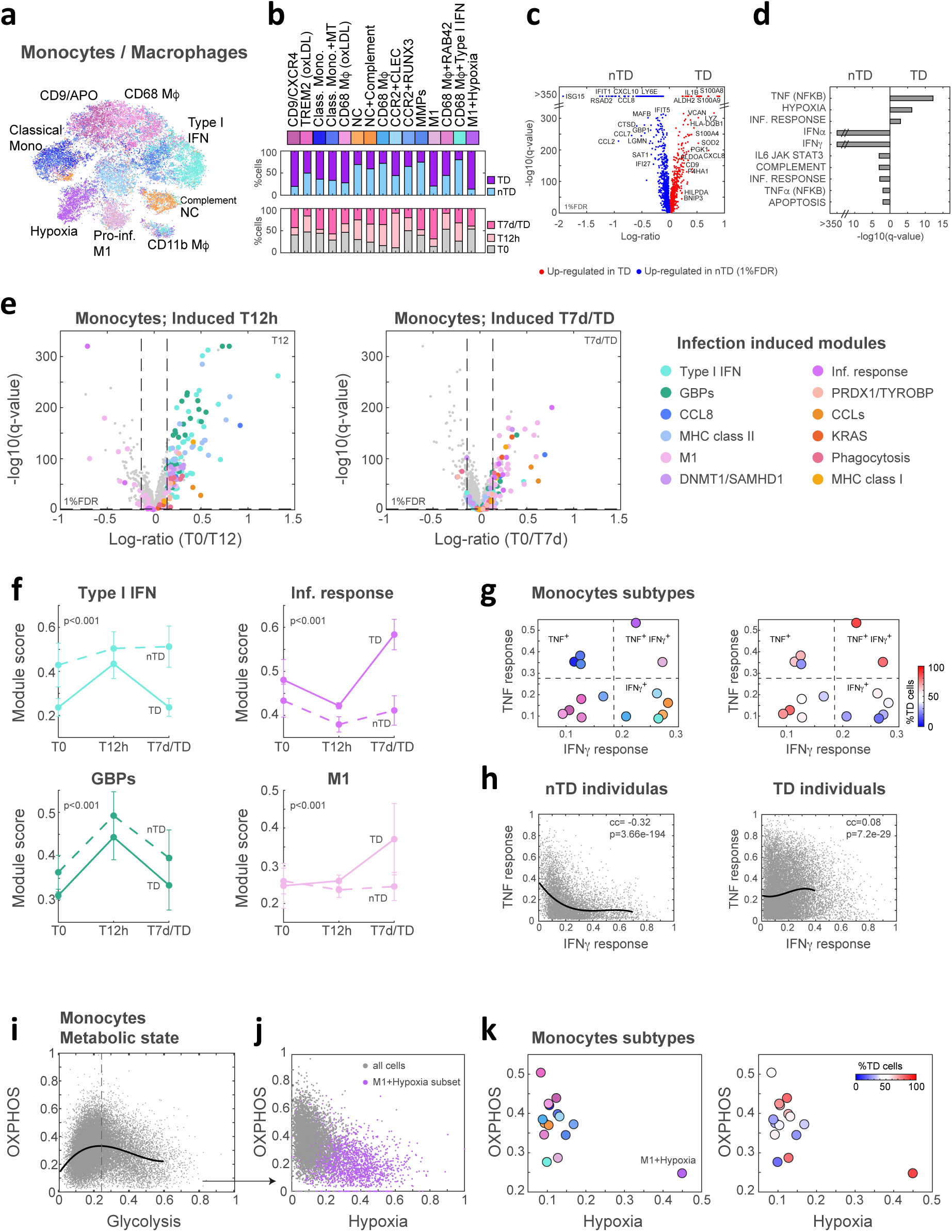
Monocyte and macrophage subset dynamics in PBMCs from the *S.* Typhi human challenge were associated with infection outcomes. (a) tSNE visualization of monocytes and macrophages from all individuals clustered to their subset level. Cells are colored by cluster identity, and subset annotations are indicated. **(b)** The composition of disease outcomes (TD and nTD) and time points (T0, T12, and T7d/TD) across monocyte and macrophage subsets show an association between specific subsets and TD or nTD individuals. **(c)** Volcano plot showing DEGs between nTD and TD individuals. The log ratio of mean expression for each gene in TD versus nTD individuals is shown on the x-axis. P-values are calculated by a two-sample t-test and q-values by FDR correction. Genes significantly up-regulated in TD individuals (red) or in nTD individuals (blue) at a 1%FDR threshold are indicated. **(d)** GSEA identifies immune and metabolic pathways significantly enriched in nTD individuals (left) and TD individuals (right). **(e)** Volcano plots displaying DEGs at early (T0 versus T12, left) and late (T0 versus TD/T7d, right) time points show different infection patterns. Genes associated with specific infection-induced modules are colored by module color (legend on the right); vertical dashed lines indicate a 1.1-fold change threshold and horizontal dashed lines indicate a 1%FDR cutoff. **(f)** Dynamics of infection-induced modules in individuals from different infection outcomes. Module scores were calculated for each individual alone across all time points. Mean scores with standard errors (SE) are shown for nTD individuals (dashed line) and TD individuals (solid line). P-values represent differences between nTD and TD dynamics, using a random model (see methods). **(g)** Expression levels for IFNγ and TNF response genes^35^ in monocyte and macrophage subsets. Dashed lines separate the four quarters of IFNγ^+/-^ and TNF^+/-^ combinations. Subsets are colored either by cluster identity (left; colors are as in (b)) or by the percentage of cells within the subset from TD individuals (right; see color bar). Co-stimulation of IFNγ and TNF (IFNγ^+^ TNF^+^) is evident only in two subsets enriched for TD individuals. **(h)** Expression of IFNγ versus TNF response in single cells separated by outcome (nTD individuals (left) and TD individuals (right)). A sliding window analysis (solid line) represents IFNγ response across TNF response levels. Correlation coefficients and p-values for TNF versus IFNγ responses are shown for each outcome group, with a positive correlation in TD individuals. **(i)** Metabolic states of monocytes and macrophages, represented as glycolysis module scores versus OXPHOS module scores in single cells. The solid line represents a sliding window of OXPHOS scores across glycolysis levels. The dashed line indicates the transition point where the derivate of the solid line equals zero (see methods). **(j)** Relationship between hypoxia and OXPHOS modules for cells with high glycolysis levels (above the dashed line in (i)). Cells from the M1+Hypoxia subset are colored in purple and located at the high hypoxic signature and low OXPHOS expression; all other cells are gray. **(k)** Metabolic states at the subset level are shown as glycolysis versus OXPHOS module scores. Subsets are colored either by cluster identity (left; colors are as in (b)) or by the percentage of cells within the subset from TD individuals (right; see color bar).

To classify underlying genes and immune pathways associated with the dynamics of disease development, we performed a pseudo-bulk analysis of the data. First, we compared the DEGs between TD and nTD individuals (**Fig. 2c and Supplementary Table 3**). Pathway enrichment analysis of these DEGs identified tumor necrosis factor (TNF) signaling, inflammatory response, and hypoxia as linked to TD outcome, while IFNα and IFNγ signaling were associated with nTD individuals (**Fig. 2d**). Second, to examine the dynamic changes induced by monocytes following *S.* Typhi challenge, we compared DEGs that exhibited significant upregulation at an early time-point following infection (T0 vs. 12h) and a late time-point (T0 vs. 7d/TD) **(Supplementary Table 4)**. We clustered these DEGs and categorized the clusters into immune-related pathways (infection-induced modules, **Supplementary Fig. 2b-c, Supplementary Table 5**). Notably, twelve hours post-infection, DEGs were related to the type I IFN and GBPs (left panel **Fig. 2e**, marked by cyan and green circles). At the late time-point (7d/TD), we observed an increase in M1 polarization genes and the inflammatory response genes (right panel of **Fig. 2e**, indicated by light pink and purple circles). Finally, to test the significance of these dynamics to disease outcome, we compared infection-induced modules between TD and nTD individuals (**Fig. 2f and Supplementary Table 5**). The type I IFN response was predominantly higher in nTD vs. TD individuals, both before challenge and at a late time point. Conversely, GBPs response was activated for both TD and nTD individuals only twelve hours post-infection. Notably, M1 polarization and the inflammatory response were observed solely in TD individuals at the time of disease diagnosis (TD).

We next focused on a monocyte subset previously identified by the co-stimulation of TNF and IFNγ pathways, which has been linked to tissue inflammation in chronic gut disease^35^. By calculating the pathway signatures for TNF and IFNγ of each monocyte subset^35^, we identified two such subsets in our data (left panel of **Fig. 2g**; TNF^+^ IFNγ^+^). These subsets, responsive to both TNF and IFNγ, were predominantly identified in individuals with typhoid disease (right panel of **Fig. 2g**). To test if this was indeed co-stimulation, we calculated the pathway signatures for TNF and IFNγ in single cells. In nTD individuals, there was a significant negative correlation within single cells in the responses, indicating that cells responded either to TNF or IFNγ signaling alone, but not to both stimuli simultaneously (left panel of **Fig. 2h**). Conversely, TD individuals exhibited activation by both stimuli together, as well as response to each stimulus alone (right panel of **Fig. 2h**; indicated by a positive correlation). Thus, the co-activation of TNF and IFNγ pathways in monocytes, linked to tissue inflammation and Crohn’s disease pathology, is observed exclusively in TD individuals and not in nTD.

We next evaluated the metabolic state of monocytes and macrophages in our data. To this end, we clustered the expression of all metabolic genes and identified clusters enriched for a specific metabolic pathway (**Supplementary Fig. 2d-e**). We identified four significant metabolic modules: glycolysis, hypoxia, oxidative phosphorylation (OXPHOS), and amino sugar and nucleotide sugar metabolism (**Supplementary Table 6**). These metabolic modules are directly linked to activation and polarization of macrophages, which entails changing their energy metabolism from OXPHOS to aerobic glycolysis^36^. To test this, we compared the OXPHOS module to the glycolysis module for each single cell and identified cells with elevated glycolysis levels and reduced OXPHOS expression (**Fig. 2i**, right to the dotted line). We further examined these cells with elevated glycolysis module and compared their OXPHOS and hypoxia modules (**Fig. 2j**). We observed two populations: one expressing the OXPHOS module and one with high levels of hypoxia module and downregulated expression of OXPHOS. These two populations resemble macrophage activation driven by aerobic glycolysis or hypoxia-driven glycolysis, respectively. Indicated in purple are the cells from the hypoxia monocyte subset (**Fig. 2b**; M1+hypoxia subset), suggesting that single cells in this cluster are indeed associated with hypoxia-driven glycolysis (**Fig. 2j**). Similarly, analysis at the subset level indicated that only the hypoxia monocyte subset had a high hypoxia score and low OXPHOS score (left panel of **Fig. 2k**). This subset was predominantly associated with TD individuals (right panel of **Fig. 2k**). Thus, we describe a unique monocyte subset that is expressing low OXPHOS, high glycolysis and high hypoxia modules, that is associated with disease outcome.

### Hypoxia gene signature in human PBMCs is associated with disease outcomes

Following the identification of immune and metabolic pathways of monocytes and macrophages specific to disease outcomes, we hypothesized that subsets of other cell types may also be associated with TD and nTD outcomes. To test this, we performed sub-clustering of the cells from each cell type. Overall, we identified 9 NK subsets, 15 T cells subsets, and 13 B cells subsets (**Fig. 3a and Supplementary Fig. 3a**). We annotated these subsets based on marker genes, including cell type markers, immune-related genes, and metabolic-related genes (**Supplementary Fig. 3a and Supplementary Table 2**). Notably, specific subsets were linked to infection outcomes, similarly to the monocytes and macrophages subsets (**Fig. 3b**). For instance, NK subsets that were enriched in individuals who developed typhoid disease expressed genes related to glycolysis, hypoxia, and proliferation. Conversely, the NK subset expressing the type I IFN response was predominantly found in individuals who did not develop the disease. Among the T cell subsets, Recent Thymic Emigrant (RTE) CD4 T cells, Th2/Th17 T cells, and T cell subset expressing hypoxia and glycolysis markers were associated with typhoid disease. Among the B cell subsets, IgM B cells (IGLC6/7+) and B cells with hypoxic markers primarily originated from individuals who developed the disease, while the type I IFN response was linked to those who did not. Collectively across all cell types, subsets that expressed the type I IFN response (IFNα) were mainly associated with individuals who did not develop typhoid disease, while subsets expressing hypoxia markers were predominantly observed in those who developed the disease. Next, we compared the DEGs between TD and nTD individuals for each cell type to further investigate the genes and pathways that are significantly different between individuals who developed the disease and those who did not (**Fig. 3c and Supplementary Table 3**). Across all cell types, TNF signaling and hypoxia were linked to disease development, while IFNα and IFNγ signaling were associated with individuals who did not develop the disease (**Fig. 3d**).

**Figure 3:**
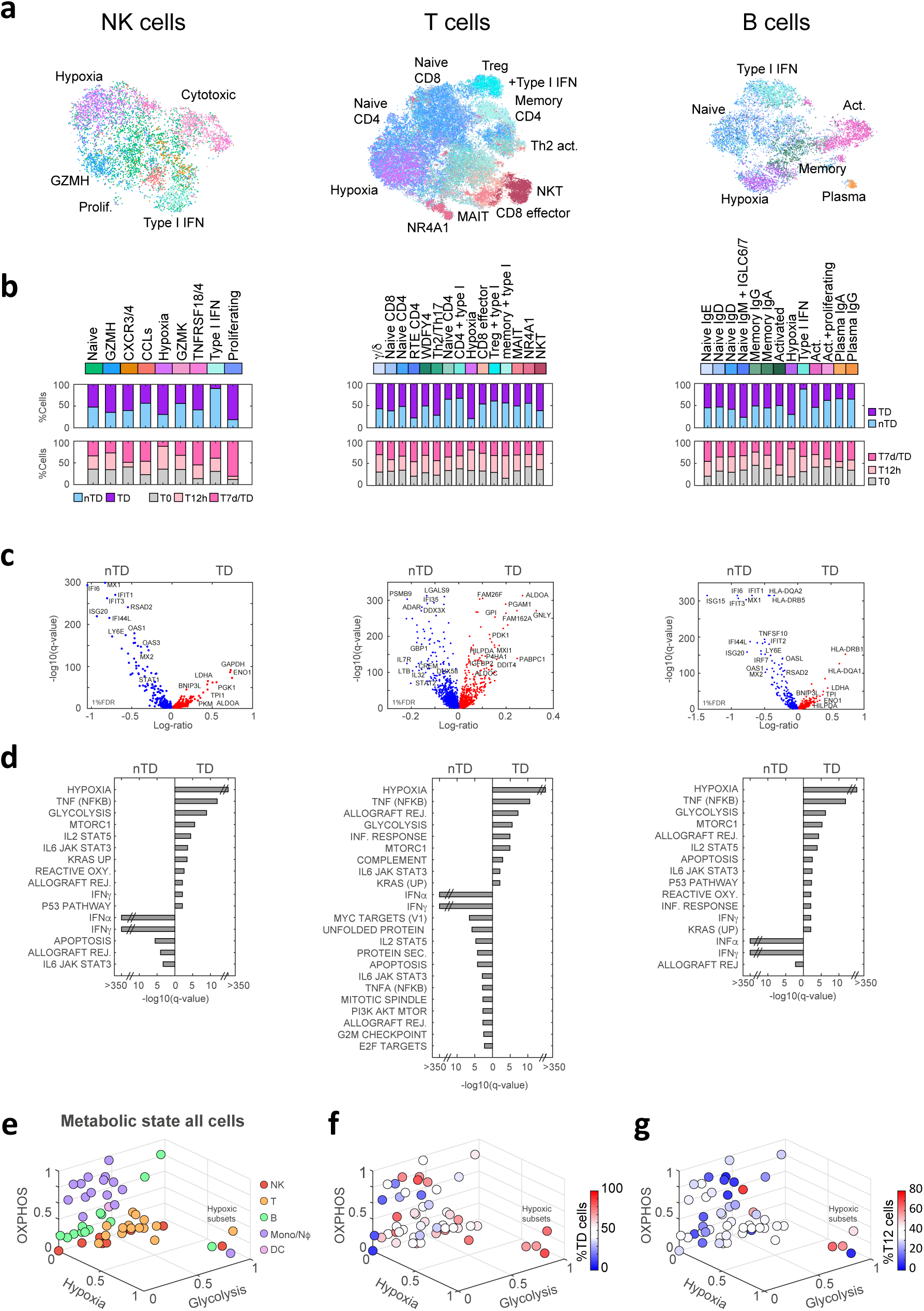
An early hypoxia gene signature in specific immune subsets from the *S.* Typhi human challenge were associated with disease outcomes. (a) tSNE visualization of NK cells, T cells, and B cells clustered to their subset level. Cells are colored by cluster identity, with subset annotations indicated. **(b)** Composition of disease outcomes (TD and nTD) and time-points (T0, T12, and T7d/TD) across NK (left), T (middle), and B cells subsets (right), showing an association of specific immune subsets with disease outcome. **(c)** Volcano plots showing DEGs between nTD and TD individuals for NK (left), T (middle), and B cells (right). The log ratio of mean expression for each gene in TD versus nTD individuals is shown on the x-axis. P-values are calculated using a two-sample t-test, and q-values are determined by FDR correction for each cell type. Genes significantly up-regulated in TD individuals (red) or nTD individuals (blue) at a 1%FDR threshold are indicated. **(d)** GSEA identifying pathways significantly enriched in nTD individuals (left) and TD individuals (right) for NK (left), T (middle), and B cells (right). **(e-g)** Metabolic states of immune subsets from all cell types are shown in a three-dimensional space representing OXPHOS, glycolysis, and hypoxia gene signatures. Subsets are colored by cell type origin in **(e)**, by the percentage of cells within the subset from TD individuals in **(f)**, and by the percentage of cells within the subset present at 12 hours post-challenge (T12) in **(g)**. A subset from each cell type shows high glycolysis and hypoxia gene signatures with low OXPHOS expression, referred to as hypoxic subsets. The hypoxic subsets are enriched in TD individuals and are evident in NK, T, and B cells already at T12.

Since the hypoxia pathway was enriched in the DEGs of all cell types, we further evaluated the metabolic states of all subsets together. We represented the metabolic state of each subset as a score within a three-dimensional space of OXPHOS, glycolysis, and hypoxia. We identified from each cell type a subset with low OXPHOS expression and high expression of glycolysis and hypoxia modules (**Fig. 3e**). Similar to the monocyte hypoxic subset, these hypoxic subsets originated from TD individuals (**Fig. 3f**). We further determined that some of these subsets were evident already as early as 12 hours post-infection (**Fig. 3g**), days before these individuals were diagnosed with typhoid disease.

Collectively, our results indicate an early hypoxia gene signature originating from unique subsets from all immune cell types, evident before the appearance of any symptoms, which is indicative of the early events at the onset of typhoid disease.

### A six-gene hypoxic signature is associated with typhoid and other infectious diseases

Having identified hypoxia-related subsets as associated with the development of typhoid disease, we generated a gene signature, capturing shared features across all of these subsets, independent of their cell-type origin (**Fig. 4a**). We identified six genes, all of them regulated by hypoxia and HIF1α: Hypoxia Inducible Lipid Droplet Associated (HILPDA) promotes lipid storage and metabolism under hypoxic conditions^37^. BCL2 Interacting Protein 3 (BNIP3) and BCL2 Interacting Protein 3-Like (BNIP3L) are both involved in promoting autophagy under hypoxic conditions^38^. They directly target mitochondria and promote the clearance of damaged mitochondria via mitophagy to prevent cellular damage. Prolyl 4-Hydroxylase Subunit Alpha 1 (P4HA1) is crucial for collagen biosynthesis and tissue remodeling under hypoxic conditions^39^. Ankyrin Repeat Domain 37 (ANKRD37) is upregulated in response to hypoxia and is implicated in cellular stress responses, cellular adaptation to hypoxia, and autophagy^40^. 1,4-Alpha-Glucan Branching Enzyme 1 (GBE1) is essential for glycan metabolism, contributing to energy storage under hypoxic conditions^41^.

**Figure 4:**
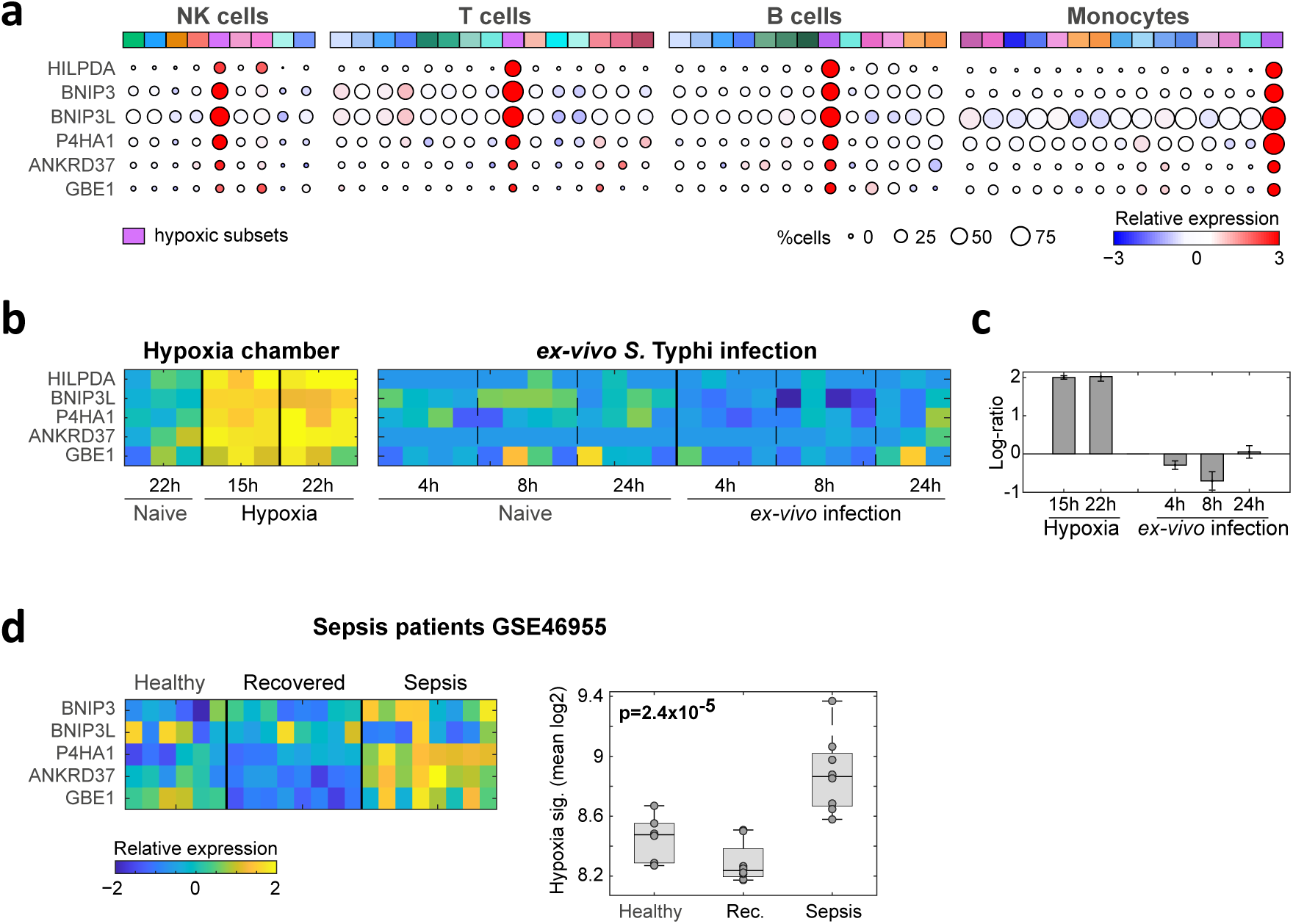
A six-gene hypoxia signature identified cells under hypoxic conditions and is associated with typhoid and other infectious disease-related pathologies. **(a)** Dot plot showing expression of six hypoxia-associated genes, specific and shared across the four hypoxic subsets (from **Fig. 3e-g**). Dot size represents the percentage of cells in a subset expressing a given gene, and dot color indicates relative expression levels (legend at the bottom). Hypoxic subsets from each cell type are indicated in purple. **(b-c)** PBMCs from a healthy donor were either subjected to hypoxic conditions (15h or 22h in a hypoxia chamber), or *ex vivo* infected with *S.* Typhi for 4h, 8h, and 24h. **(b)** Heat map showing the expression levels of the six-gene hypoxia signature in PBMCs under hypoxic conditions (left) or following *ex vivo* infection (right). **(c)** Quantification of the hypoxia signature fold-change relative to matched naïve samples for both hypoxic conditions and *ex vivo* infection. The signature is significantly elevated only under hypoxic conditions. Mean signature and SE values are shown. **(d)** Heat map of expression levels of the six-gene hypoxia signature in a dataset of isolated blood monocytes from sepsis patients during active sepsis, post-recovery, and from healthy controls. Box plot comparing six-gene hypoxia signature across conditions indicates significant induction of the signature only during sepsis; a p-value was calculated using one-way ANOVA (indicated in the figure). The color bar below the heatmap indicates relative expression levels corresponding to (b) and (d).

The six-gene hypoxia-related signature may be driven either by bona-fide hypoxic conditions or, alternatively, HIF1α activation in response to infection through lipopolysaccharides (LPS) and Toll-like receptor 4 (TLR4) signaling^42^. To test this, we incubated PBMCs from a healthy donor in a hypoxia chamber for 22 hours to induce low-oxygen conditions, or infected PBMCs *ex-vivo* with *S.* Typhi for 24 hours under normoxic conditions. Using bulk RNA-seq analysis of these samples, we curated HIF1α target genes that were induced by both hypoxia and *ex-vivo* infection, solely by hypoxia or solely by infection (**Supplementary Fig. 4a-b and Supplementary Table 7**). Our six-gene signature was induced exclusively by hypoxia (**Fig. 4b-c**). We further analyzed the expression of additional hypoxia gene signatures, to establish our hypoxia-related subsets. Specifically, we tested three hypoxia-related gene signatures from cancer patient databases, where hypoxia conditions are well-documented^43–45^. All of these signatures were highly expressed across our hypoxic subsets - NK, T, B cells, and monocytes (**Supplementary Fig. 4c**). Importantly, these signatures were distinct from each other and from our six-genes signature (**Supplementary Fig. 4d**), further supporting hypoxia-related signatures of these subsets.

We next studied the expression of our six-gene hypoxia signature in other human infectious disease-related pathologies known to involve hypoxic conditions. We first analyzed a public dataset of Gram-negative sepsis patients, where tissue hypoxia is a prevalent pathophysiological feature^46^. This dataset included expression data of blood monocytes from patients during sepsis, after recovery, and healthy controls. Our six-gene hypoxic signature was significantly elevated in septic patients compared to post-recovery levels and healthy controls (**Fig 4d**). Next, we evaluated our signature in a dataset of Crohn’s disease patients, which included gut samples from patients and healthy controls^47,48^. The signature was detected exclusively in a subset of macrophages present only in Crohn’s disease patients, particularly in cases of severe disease, and not in normal controls (**Supplementary Fig. 4e-i**). Last, we examined a public dataset of patients with cutaneous leishmaniasis, as these lesions are known to promote hypoxic microenvironment^49^. We analyzed the expression of genes significantly elevated in hypoxic lesions of cutaneous leishmaniasis compared to non-hypoxic lesions^50^. Notably, genes associated with hypoxic lesions were enriched in the hypoxic subset of the monocytes in our dataset (**Supplementary Fig. 4j**). Collectively, our six-gene hypoxic signature is observed in a range of pathologies associated with infection and tissue hypoxia.

### Transcriptional changes of immune cells in the gut in a mouse model of typhoid fever

To gain mechanistic understanding of the immune subsets we observed in the human challenge model and establish a link to tissue-specific immune responses at the site of infection, we utilized a mouse typhoid model. The typhoid fever mouse model has long been useful for the valuable insights it has offered into the invasion and systemic dissemination of *S.* Typhi infection in humans^2^. In this model, *Nramp1-*sensitive mice are orally infected with *S.* Typhimurium in the absence of streptomycin pretreatment^4^. After oral infection, ingested *S.* Typhimurium enters the small intestine and invades the Peyer’s Patches (PP) located in the terminal ileum, after which it disseminates to the blood, lymph nodes, and systemic sites^6,7^. To assess the transcriptional changes occurring during infection, we orally infected C57Bl6/J mice with *S.* Typhimurium strain SL1344 and conducted bulk RNA sequencing (RNA-seq) on selected tissues, including the ileum (small intestine), cecum (large intestine), PP, and the spleen, as a key site for systemic infection. We analyzed RNA-seq data before infection and at two time points post-infection: 24 hours and 72 hours (**Fig. 5a**). Variation among samples was mainly attributed to the differences between the tissues, as indicated on the Principal Component Analysis (PCA; **Fig. 5b**, **Supplementary Fig. 5a-b, and Supplementary Table 8**). When analyzing each tissue independently, we found that 72 hours post-infection the PP exhibited the most pronounced changes, with 200 differentially expressed genes (DEGs), compared with 25 DEGs in the spleen and only one DEG in the ileum and cecum (10% FDR; **Fig. 5c-e**, **Supplementary Fig. 5c-d and Supplementary Table 9**). Gene Set Enrichment Analysis (GSEA) revealed several significantly altered immune and metabolic pathways 72 hours post-infection, including TNF signaling, IFNγ and IFNα pathways, and inflammatory response (**Fig. 5f and Supplementary Fig. 5e**). Importantly, in the PP we also observed a hypoxia-enriched signature reminiscent of the hypoxia signature in the human challenge model (**Fig. 5f-g**).

**Figure 5:**
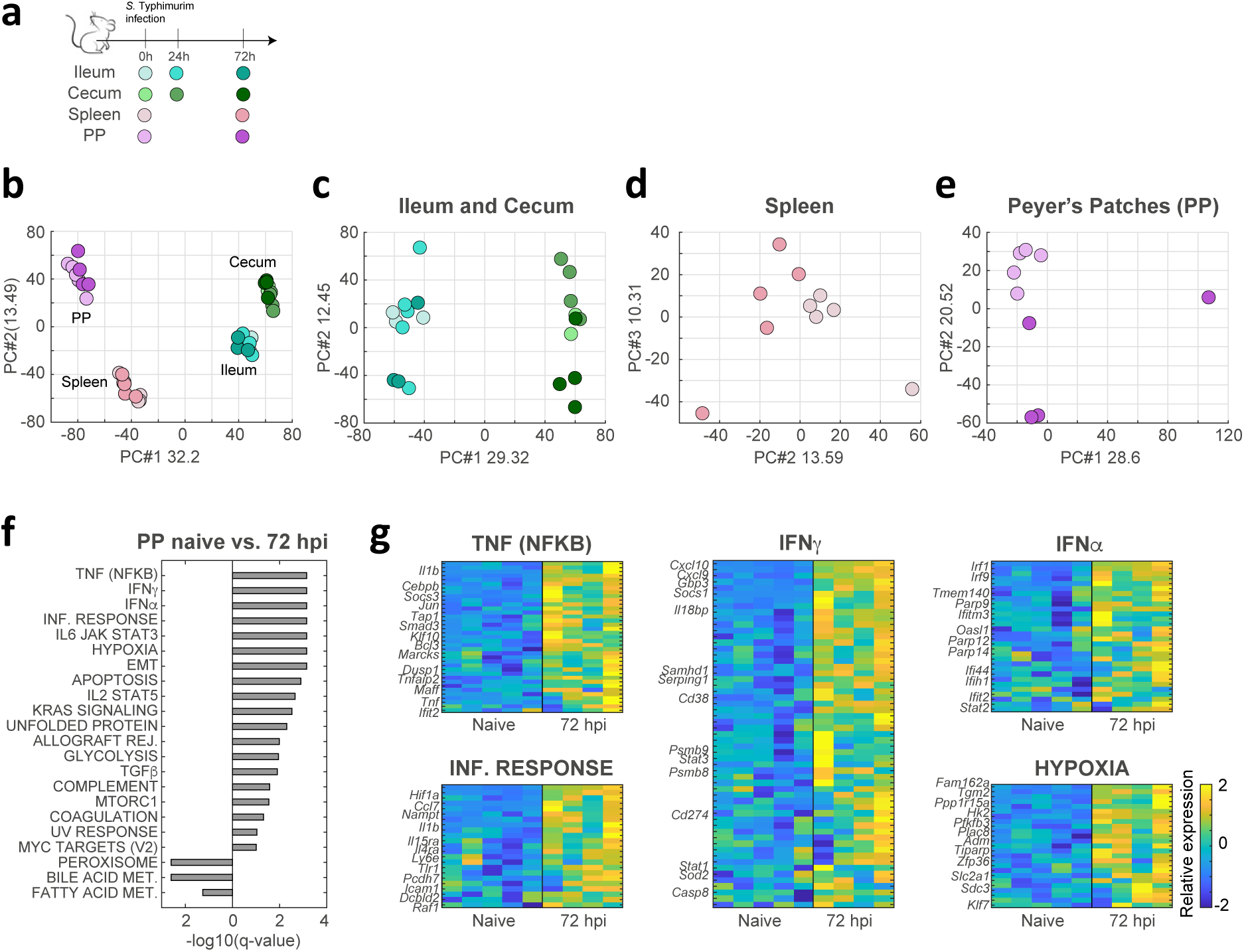
Transcriptional changes of immune cells in the gut in a mouse model of typhoid fever. **(a)** Overview of the experiment: mice were orally infected with *S.* Typhimurium, without streptomycin pretreatment. Immune enriched samples were collected from ileum, cecum, spleen, and Peyer’s Patches (PP) before infection and 24 or 72 hours post-infection (hpi) and analyzed by bulk RNA-seq. **(b-e)** Principal component analysis (PCA) projection of the bulk RNA-seq data onto the space of the two leading principal components for all samples together **(b)**, ileum and cecum samples alone **(c)**, spleen samples **(d)**, and PP samples **(e)**, showing tissue-specific separation independent of infection, and infection-induced separation within the spleen and most pronounced in the PP. Color coding for tissues and time points corresponds to (a). **(f)** Gene Set Enrichment Analysis (GSEA) comparing *S.* Typhimurium infected PP at 72 hpi versus naïve PP identifies up-regulated (right) and down-regulated (left) immune and metabolic pathways in infected PP. **(g)** Heatmaps of the top significant pathways from (f), showing the expression levels of the corresponding genes in naïve versus infected PP. The color bar indicates relative expression levels, and representative genes for each pathway are highlighted.

As we found the hypoxia signature only in the PP, we next characterized cell type-specific immune responses and immune population changes to reveal immune subsets that specifically express hypoxia signatures. PP samples were enriched for immune cells (using Ficoll), before and 72 hours after infection, and analyzed by scRNA-seq. In total, we sequenced 2525 cells from naïve and infected mice, identifying five immune cell types (CD4 T cells (*Cd3g* and *Cd4*), cytotoxic CD8 T cells (*Cd3g*, *Cd8a*, and *Xcl1*), B cells (*Cd19* and *Cd79b*), Germinal center (GC) B cells (*Cd19* and *Aicda*), and macrophages (*Lyz2* and *Csfr1*)) as well as three non-immune cell types (endothelial cells (*Pecam1*), Paneth cells (*Defa21/22/24*), and epithelial cells (*Epcam* and *Lgals2*) (**Fig. 6a-d**). At 72 hours post-infection, we observed a reduction in GC B cells (**Fig. 6c**), consistent with previous findings indicating that following bacterial infection, GC collapses due to hypoxia and impaired cellular respiration^51^. Additionally, we noted an infiltration of macrophages into the PP after infection (**Fig. 6c**). To evaluate cell type-specific infection responses within the PP, we analyzed in each cell type the immune and metabolic responses observed in the bulk RNA-seq data (from **Fig. 5g**). Notably, our results identified that the TNF and inflammatory response were primarily driven by infiltrating macrophages, while responses to IFNγ and IFNα were noted across all cell types (**supplementary Fig. 6a-c**). We found that hypoxia gene signature was primarily elevated in infiltrating macrophages but was also evident in other cell types, including B cells (**Supplementary Fig. 6c**). Finally, we examined our six-genes hypoxia signature from the human challenge model (**Fig. 4a**) in the mouse PP scRNA-seq dataset. Notably, the hypoxic signature was evident in a macrophages subset that infiltrated the PP 72 hours post-infection (**Fig. 6e**), and to a lesser extent also evident in other cell types (**Supplementary Fig. 6d**).

**Figure 6:**
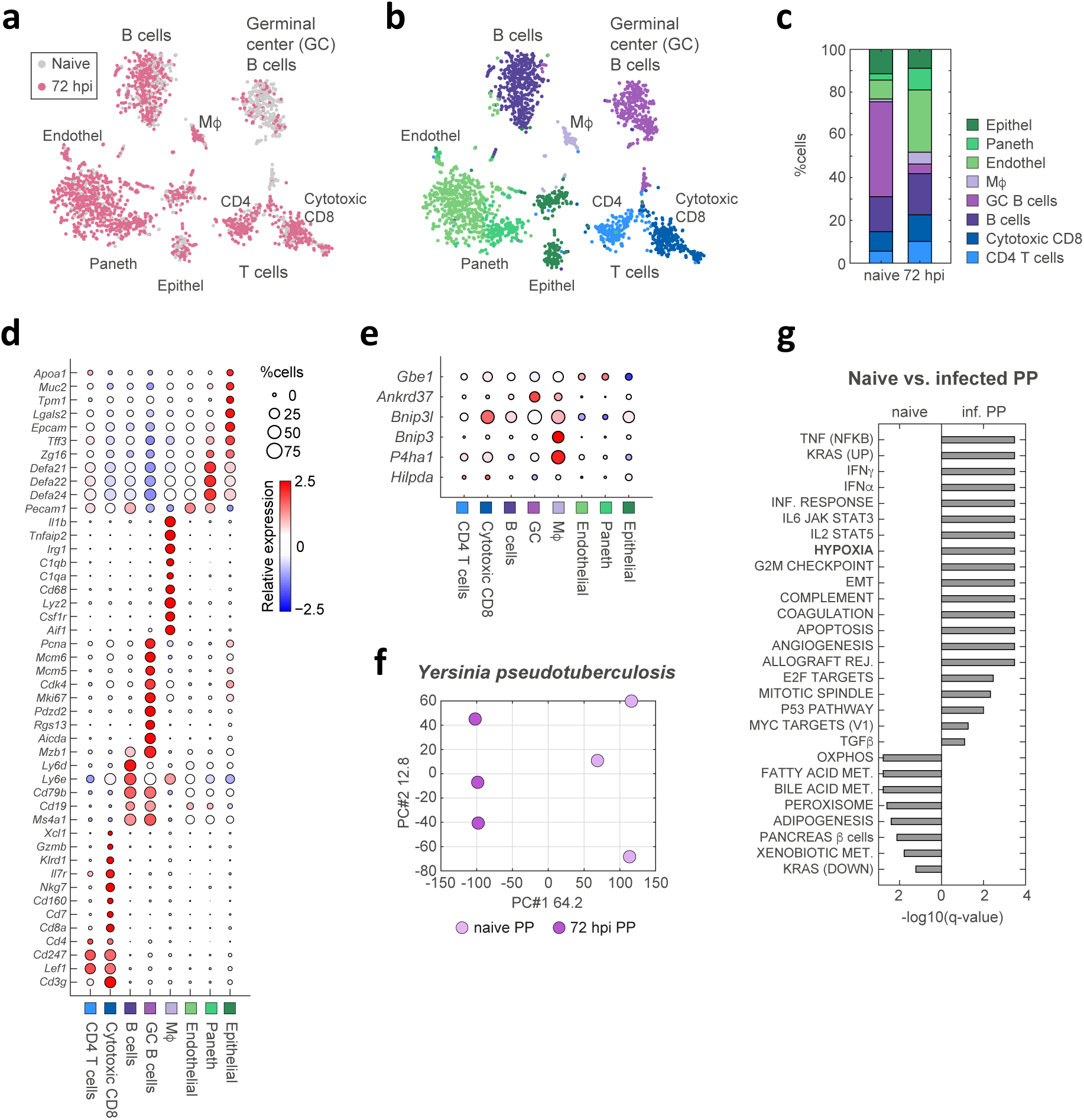
scRNA-seq analysis of immune cells in the PP in a mouse model of typhoid fever. (a-b) tSNE visualization of scRNA-seq data from naïve and infected PP at 72 hpi. Cells are colored by infection status (naïve or infected) in **(a)**, and by cell clusters in **(b)**, with cluster annotations indicating cell types. **(c)** Comparison of cell-type composition between naïve and infected PP shows macrophage (M<) infiltration and a reduction in Germinal center (GC) B cells in infected PP. Colors correspond to clusters in (b), with cell-type annotations shown in the legend on the right. **(d)** Dot plot showing expression levels of marker genes across clusters for each cell type. The size of each dot represents the percentage of cells in a cluster expressing a given gene, and the color indicates relative expression levels (legend on the right). **(e)** Dot plot showing expression levels of the six-gene hypoxia signature in immune cells from the PP of infected mice. **(f-g)** Analysis of public bulk RNA-seq data from the PP of naïve and *Yersinia pseudotuberculosis*-infected mice (72 hpi)^18^. **(f)** PCA projection based on the two leading principal components, showing separation between PP of naïve and *Yersinia* infected mice. **(g)** GSEA comparing gene expression in *Yersinia* infected versus naïve PP identifies up-regulated (right) and down-regulated (left) pathways. The hypoxia pathway is highlighted and significantly up-regulated in *Yersinia*-infected PP relative to naïve PP.

We also examined an RNA-seq dataset of PP infected with *Yersinia pseudotuberculosis*^18^, a Gram-negative, food-borne pathogen that similarly invades and colonizes the PP. GSEA analysis of the DEGs between naive and infected PP 72 hours post-infection identified an elevation in the hypoxia pathway in infected PP (**Fig. 6f-g**), similar to what we observed in the mouse typhoid model.

Collectively, transcriptional analysis of mouse infection models indicated that hypoxia-related signatures in specific immune subsets are associated with colonization and invasion of pathogens to the PP.

### The hypoxic signature is present in the PP of infected mice and depends on the pathogen’s invasion and colonization

To functionally test if invasion of the pathogen into the PP is required for hypoxia-related gene signature, we infected orally C57Bl6/J mice with either wild-type IR715 *S.* Typhimurium strain or a triple mutant IR715 strain (*invA, fliC, fljB* mutant). The triple mutant lacks the flagellin genes *fliC* and *fljB* and is deficient in the invasion-associated type III secretion system (T3SS-1) gene *invA,* rendering this strain unable to invade and colonize the mouse PP^52^. We verified that this mutant strain is impaired in its ability to invade and colonize the PP in the mouse typhoid model, by plating the PP of wild-type and triple mutant infected mice for colony forming units (CFU) (**Fig. 7a**). Bulk RNA-seq analysis of PP samples from mice infected with either the wild-type or triple mutant strain identified distinct expression patterns compared to naïve mice (**Fig. 7b and Supplementary Fig. 7a**). GSEA of the DEGs during wild-type infection or triple mutant infection showed that the hypoxia pathway was elevated only in infection of wild-type *S.* Typhimurium and not in triple mutant infected mice (**Fig. 7c**). These results suggest that the hypoxic signature of immune cells depends on pathogen invasion and colonization in the PP.

**Figure 7:**
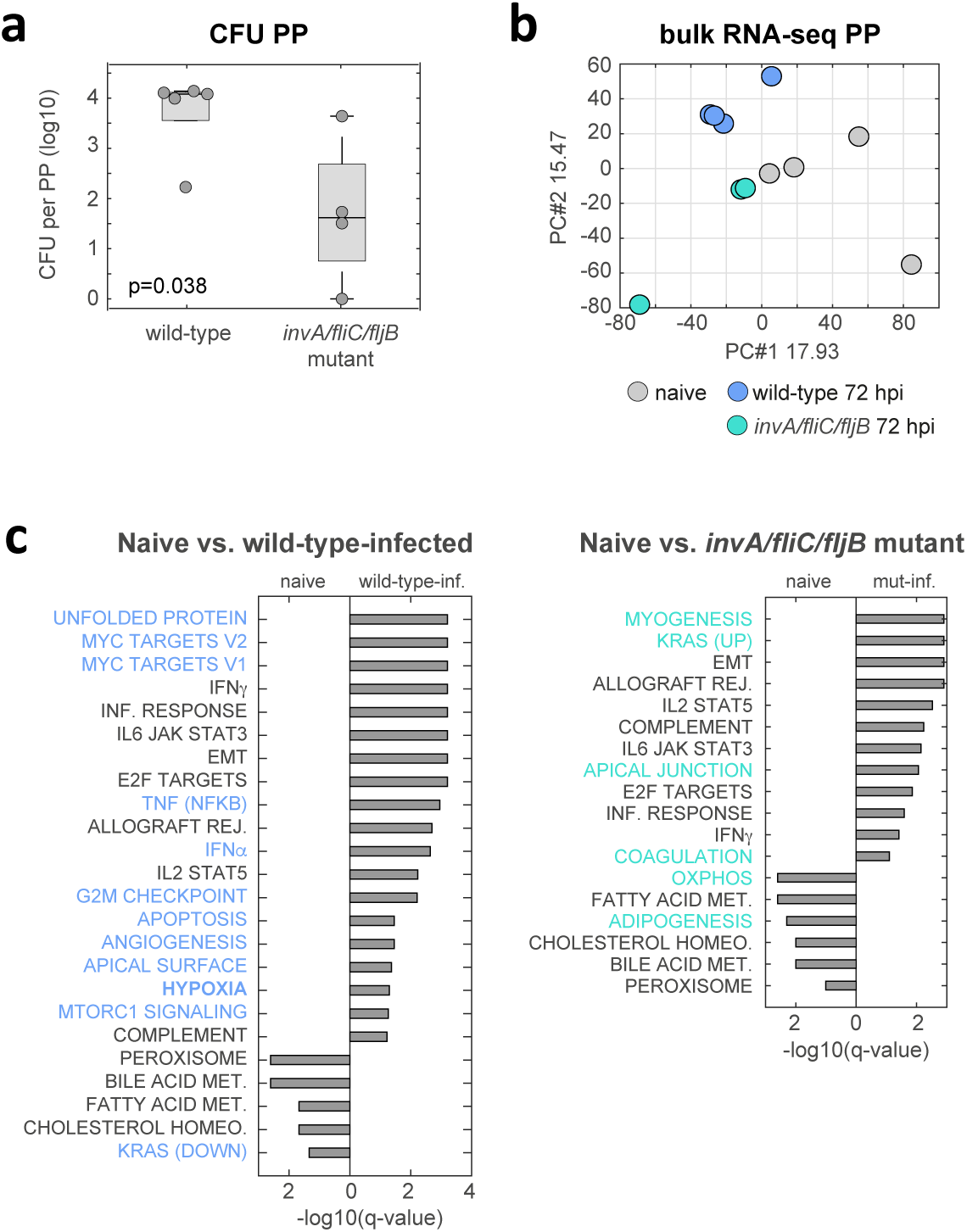
Hypoxia signature was dependent on pathogen’s invasion and colonization of the PP in the mouse typhoid disease model. (a-c) Mice were orally infected with either wild-type *S.* Typhimurium or a triple mutant strain (*invA/fliC/fljB* mutant). **(a)** Bacterial burden in the PP of wild-type and triple mutant strain infected mice was quantified by colony forming units (CFU). **(b)** PCA projection based on the two leading principal components, showing separation between PP of naïve, wild-type infected, and triple mutant-infected mice. **(c)** GSEA comparing wild-type infected PP at 72 hpi versus naïve PP (left), and triple mutant-infected PP versus naïve PP (right) identifies strain-specific pathways (highlighted in blue for wild-type strain and in turquoise for mutant strain). The hypoxia pathway is significantly up-regulated only in PP infected with the wild-type strain, but not in triple mutant infected PP.

In summary, our results demonstrate that infection in a mouse model of typhoid disease induced a hypoxic signature in the PP, contingent on pathogen invasion and colonization. Additionally, this hypoxic signature appears early after *S.* Typhi infection in a human challenge model, and it is associated exclusively with individuals who later develop bacteremia and systemic typhoid disease.

## Discussion

In this study, we performed an integrated analysis of an animal model and a human challenge model of infection. We profiled immune responses to infection in the gut in a mouse model of typhoid disease, and of peripheral immune cells from participants in the *S.* Typhi human challenge model. The mouse typhoid model simulates *S*. Typhi infection in humans, with bacteria invading the mucosal barrier into the PP and systemic sites without triggering overt gut inflammation^5,6^. Accordingly, our findings indicate the PP as the primary site with transcriptional and immune subset changes in response to infection. Within the PP, we identified a hypoxia-related subset of infiltrating macrophages after infection. The recruitment of this subset was dependent on bacterial invasion and colonization of the PP, as mice infected with a mutant strain impaired in PP invasion did not exhibit this hypoxia-related macrophage signature. Moreover, alongside the infiltration of macrophages with the hypoxic signature, we observed a reduction in GC B cells, as previously reported^51^. In humans, we found a similar hypoxia-related signature in immune subsets enriched in individuals who developed typhoid disease and not in those who remained asymptomatic. We verified that this signature was induced by hypoxic conditions rather than HIF1α activation through LPS and TLR4 signaling^42^, as immune cells incubated in a hypoxic chamber elevated this signature, whereas *ex vivo* infected cells did not. Interestingly, hypoxic immune subsets were detectable in humans several days before disease onset, as early as 12 hours post-challenge. We also measured the hypoxia-related signature in other infectious disease-related pathologies, such as sepsis and Crohn’s disease patients, suggesting broader implications.

Hypoxia has long been recognized as an important feature of gut physiology. The gut mucosal tissue maintains a hypoxic environment under steady-state conditions, required for normal barrier function, healthy microbiome, and mucosal immunity^53^ (e.g., GC B cell function and antibody class switching^54^). During infection, especially in the context of *S.* Typhimurium infection in the setting of antibiotic-induced gut dysbiosis^55^, gastrointestinal inflammation disrupts the oxygen balance^56^, leading to damage of epithelial cells^57,58^, disruption of the intestinal barrier^59^, and alterations to the microbiota architecture^11,60,61^. However, the role of hypoxia in the context of gut mucosal immunity, particularly in the absence of overt inflammation and tissue damage, as observed in the typhoid model, remains poorly understood. Hypoxia during immune challenge arises from increased metabolic demand of inflamed resident cells, infiltration of activated immune cells, and multiplication of intracellular pathogens, all of which deprive local immune cells of oxygen^62,63^. Our findings of infiltrating macrophages with a hypoxia-related signature indicate a role for hypoxia in the PP in the typhoid mouse model. In a recent study, hypoxia in the PP of mice during *S.* Typhimurium infection was shown to drive GC B cell death, GC collapse, and disruption of antibody-mediated immune responses^51^. Blocking secretion of TNF by macrophages in the PP restored GC function^51^. Future studies may further delineate the function of our TNF-secreting M1-hypoxia macrophage subset in driving hypoxia and GC collapse in the PP.

Our findings of unique subsets with a hypoxia-related signature in human peripheral blood raise the hypothesis that they may originate from subsets with a hypoxia signature in the PP, as indicated by our mouse infection model. This hypothesis is supported by observations of immune cell trafficking from the PP to the circulation under steady-state conditions, which is considerably accelerated during inflammation^64^. However, given the blood is not a hypoxic environment, persistence of the hypoxia-induced transcriptional changes in these cells, even after they leave the hypoxic environment of the gut to the circulation, requires further exploration. One hypothesis is that the hypoxia-related signature is associated with epigenetic changes during hypoxia in the PP, which then migrate to the blood of infected individuals. The core six genes in our hypoxic signature are related to cellular damage processes. BNIP3 and BNIP3L, which promote autophagy under hypoxic conditions^38^, regulate DNA damage and are linked to the induction of senescence^65^. Additionally, BNIP3, an outer mitochondrial transmembrane protein, regulates the expression of p16, a key senescence regulator, and its induction is required for DNA damage in the context of senescence^66^. Last, the cytokine expression observed in our hypoxia-related macrophage subset in the gut of the typhoid mouse model and in the periphery of humans challenged with *S.* Typhi resembles the overt production of inflammatory cytokines and chemokines collectively known as senescence-associated secretory phenotype (SASP)^67^. Future studies may delineate the role of DNA damage in imprinting the hypoxia signature in circulating immune subsets.

Our study underscores the value of integrating animal models with human studies to better understand the mechanism underlying infection outcomes. On the one hand, PBMCs have long been recognized as invaluable for studying the complexity of the human immune system in both health and disease^68,69^. Blood contains all major immune cell lineages and is easily accessible, making blood-derived immune cells a widely used resource for investigating how various pathogens influence immune responses^31–33,70–72^. However, PBMCs alone are limited to circulating cells, that may not represent immune responses occurring at the site of the infection. Furthermore, given that human challenge studies can be performed only on limited numbers of individuals, the analysis of immune correlates to infection outcomes is underpowered. On the other hand, mouse models can provide an understanding of immune responses at the local tissue, which is critical for a comprehensive understanding of host-pathogen interactions^73^. However, the immune responses of mice may not translate directly to human infection^74,75^. Furthermore, as all infected animals develop disease, mouse models do not provide insight into immune responses linked to infection outcomes. Our combined approach allowed high-resolution mapping of immune responses at single-cell resolution, overcoming the limitations of each model, and bridging the gap between immune responses at the site of infection in mice and immune correlates of infection outcomes in humans.

Together, our results suggest that after oral *S.* Typhi infection, the invasion and colonization of the pathogen in the PP induces hypoxia-related immune subsets, detectable early in the blood of susceptible individuals. A long-standing controversy in the field is what underlies inadequate antibody responses during bacterial gut infection and impedes vaccine development^76^. Thus, our hypoxia-related signature may serve not only as a potent biological marker of early gut invasion, but may offer a mechanism to prevent damage to the PP and improve protective antibody responses that can lead to a rational design of a new generation of vaccines for bacterial pathogens.

## Methods

### Human typhoid infection model

PBMCs samples in the current study are part of the samples collected in the *in vivo* human challenge model from study OVG2014/08 (VAST)^21^. In the VAST study, an observer and participant-masked, randomized, controlled, phase 2b study was done at the Centre for Clinical Vaccinology and Tropical Medicine (Churchill Hospital, Oxford, UK). Healthy volunteers were recruited at the ages of 18 and 60 years, with no previous history of typhoid vaccination, infection, or prolonged residency in a typhoid-endemic region. All volunteers underwent extensive medical screening (see Jin et al.^21^ for full inclusion and exclusion criteria). Written informed consent was obtained from all volunteers before enrolment. The study protocol was approved by the sponsor (University of Oxford), the South Central Oxford A Ethics Committee (14/SC/1427), and the Medicines and Healthcare Products Regulatory Agency (Eudract 2014-002978-36). The study was done in accordance with the principles of the Declaration of Helsinki and the International Council for Harmonisation Good Clinical Practice guidelines. Participants were randomly assigned (1:1:1) to receive a single parenteral dose of Vi-tetanus toxoid conjugate (Vi-TT; Typbar-TCV, Bharat Biotech, Hyderabad, India), Vi-polysaccharide (Vi-PS; TYPHIM Vi, Sanofi Pasteur, Lyon, France), or control meningococcal ACWY-CRM conjugate vaccine (control; MENVEO, GlaxoSmithKline, Sovicille, Italy). Individuals used in this study are all from the control group of the VAST study, who got the meningococcal ACWY-CRM conjugate vaccine. Following vaccination, participants completed an online diary for 7 days to monitor local and systemic symptoms, and were assessed in clinic on days 1, 3, 7, and 10. About one month post-vaccination, participants were challenged orally with 1–5 × 10^4^ colony forming units (CFUs) of *Salmonella* Typhi Quailes strain (a wild-type strain originally isolated from a chronic carrier in Baltimore, MD, USA). Immediately before challenge administration, participants ingested 120 mL of sodium bicarbonate solution to neutralise gastric acid^19^. Following the challenge, participants were seen daily for vital sign measurement, blood collection, and general assessment in an outpatient clinic for a 2 week period and completed an online diary with twice daily self-reported temperature measurements for 21 days, covering the 2-week challenge period and an additional 7 days after challenge to monitor antibiotic tolerability and symptom resolution. Typhoid fever was diagnosed if pre-determined criteria were met: a positive blood culture with *S* Typhi collected more than 72 h post-challenge or a fever of 38°C or higher persisting for 12 h or longer (participants were denied access to antipyretics before diagnosis). Diagnosed participants commenced a 2-week course of antibiotics, either ciprofloxacin 500 mg twice daily or azithromycin 500 mg daily, and attended five follow-up appointments to monitor disease resolution. Participants diagnosed on the basis of positive blood culture were commenced on treatment when blood cultures flagged positive with Gram-negative bacilli (with confirmatory identification of *S.* Typhi done after treatment commencement). Participants who did not develop typhoid fever despite oral challenge (undiagnosed) were treated with antibiotics at the end of the challenge period (day 14).

In the current study, we have used samples from 6 individuals enrolled in the control group (received meningococcal ACWY-CRM conjugate vaccine). Three of them were categorized as developed typhoid fever (TD), and three did not develop the disease throughout the challenge period (nTD). For each of the 6 individuals, 3 PBMC samples were tested in the current study: before the challenge, 12h after the challenge, and the last sample was taken when symptoms appeared (for the TD individuals) or on day 7 after the challenge (for nTD individuals), the median time point for disease diagnosis. In total, 18 PBMC samples were used in this study from the human typhoid infection model.

### PBMCs from healthy donors

Lukocyte-enriched fractions from healthy donors were obtained from the blood bank (MDA, Israel). Blood samples sourced from the blood bank are anonymized, with no donor-identifiable information, to ensure confidentiality and compliance with ethical standards. PBMCs were extracted, aliquoted to 20* 10^6 cells/ tube, and frozen as described in Hen-Avivi and Avraham 2022^77^. For each experiment, PBMCs from one donor were defrosted, counted with viability staining (ViaStain™ AOPI Staining Solution, Revvity CS2-0106-5ml) resuspended in RPMI+ media (RPMI 1640 with L-Glutamine supplemented with 10% heat-inactivated fetal bovine serum and 1mM sodium pyruvate), plated on U-bottom 96-well-untreated plates (5* 10^5 cells/well unless mentioned otherwise) and incubated at 37°c and 5% CO_2_ over-night for resting before further treatment.

### Hypoxic conditions for PBMCs

PBMCs from one healthy donor were defrosted and treated as described in ‘PBMCs from healthy donors’ in the method section. The day after, PBMCs were either kept in normoxia (37°c incubator with 5% CO_2_) for 22 hours or transferred to a hypoxia chamber (2% O_2_) for 15 or 22 hours, followed by RNA extraction.

### Salmonella Typhi growth and PBMC infection

*Salmonella* Typhi Quailes strain was inoculated into LB media and grown for 14 hours at 37°c to stationary phase with tightly closed tubes on a roller. After 14 hours, the bacteria were centrifuged (1min 11,000 RPM), the media was discarded, and the bacteria were resuspended with DPBS, with calcium and magnesium (Sartorius, 02-020-1A, from now on DPBS). OD 600 was measured, and the bacteria were diluted to OD 600 0.1. PBMCs to be infected were defrosted and treated as described in ‘PBMCs from healthy donors’ in the method section one day before the infection. For each condition, 4 PBMC wells were infected (or uninfected as controls) from the same PBMCs donor and used as replicates. For well, 5ul of the diluted bacteria stock (equal to MOI of 1, note that MOI refers to the number of total PBMCs in the well and not only to monocytes) or DPBS for controls were added to the PBMCs, the PBMCs were gently mixed by pipetting and incubated for 30 minutes for internalization in a 37°c incubator with 5% CO_2_. After 30 minutes, the PBMCs were centrifuged (500g for 5 min in RT), the medium was discarded, and the cells were washed with RPMI+ media supplemented with gentamycin (50 ug/ml) to eliminate *Salmonella* that were not internalized. The cells were centrifuged again (500g for 5 min in RT), the media was discarded, and the cells were again resuspended with RPMI+ supplemented with gentamycin (50 ug/ml). The cells were then incubated for 4, 8, and 24h in a 37°c incubator with 5% CO_2_ before being used for RNA preparation. The diluted bacterial stock used for infection was validated for the desired concentration by plating serial dilutions on LB agar plates followed by overnight incubation in a 37°c incubator and counting the grown colonies.

### Salmonella Typhimurium SL1344, IR715 and FR90 growth for mice infection

*Salmonella* Typhimurium strains were inoculated into LB media supplemented with 0.3M NaCl and antibiotics (see antibiotics per stain below), and grown for 16 hours in 37°c on a roller to provide consistent agitation. After 16 hours, the bacterial culture was diluted 1:20 with fresh media (LB media supplemented with 0.3M NaCl+ antibiotics) and grown for 4 hours. The culture was then centrifuged (1min 13,000 RPM), the media was discarded, and the bacteria were resuspended with DPBS. OD 600 was measured, and the bacteria were diluted to OD equal to 1*10^9^ bacteria in 100 ul (equal to 1*10^10^ bacteria/ml) in DPBS. The diluted bacterial stock used for infection was validated for the desired concentration by plating serial dilutions on LB agar plates followed by overnight incubation in a 37°c incubator and counting the grown colonies. SL1344 is naturally resistant to streptomycin, and 100 μg/ml streptomycin was added to the media when it was grown. *Salmonella* Typhimurium strains IR715 (nalidixic acid derivative of ATCC 14028S)^78^ and FR90 (IR715 *invA fliC fljB*, triple mutant, nalidixic acid, kanamycin, and ampicillin-resistant)^52^ were kindly provided by Andreas Baumler lab. IR715 was grown in the presence of 50 µg/ml nalidixic acid, and FR90 IR715 was grown in LB+ 50 µg/ml nalidixic acid+ 50 µg/ml kanamycin kanamycin + 100 µg/mL ampicillin.

### Mice gavage infection

Female C57BL/6 mice, 7-9 weeks old, were purchased from ENVIGO, housed at the Weizmann Institute of Science pathogen-free facility, and provided with standard food and water without restriction. Water and food were withdrawn for 4h before the mice were orally infected with 1*10^9^ CFU of Salmonella Typhimurium strain SL1344 (in 100μl suspension in DPBS) or treated with sterile DPBS (control). Ten to fifteen min before infection, mice were orally treated with 100μl of 1.3% W/V sodium bicarbonate or sterile DPBS (control) to reduce acidity in the gut, a procedure that was shown to increase infection^79,80^. Thereafter, drinking water ad libitum was offered immediately, and food was provided 2h post-infection. At the indicated times, mice were sacrificed by cervical dislocation, and the desired tissue (ileum, caecum, Peyer’s patches (PP) and/ or spleen) were removed for analysis.

### Mice tissue extraction

#### Isolation of immune cells from mice ileum and cecum

ileum and cecum from 3 naïve mice, 5 infected mice for 24h and 4 mice infected for 72h were collected as follows: the protocol was done based on Gross-Vered et al. 2020^81^. Five cm of distal ileum and cecum were collected and kept on ice-cold DPBS. Extra-intestinal fat tissue and PP were carefully removed, and organs were then flushed of their luminal content with cold DPBS, opened longitudinally, and cut into 0.5 cm pieces. Epithelial cells were removed by washing in cold DPBS using handshaking and passing through a 70 mm cell strainer twice. Epithelial cells were removed by incubation with 2mM EDTA + 20mM Hepes in DPBS without calcium and magnesium (Sartorius, 02-023-1A, from now on DPBS -/-) at 37°C incubator with shaking at 250 rpm (25 min incubation for ileum extraction, 30 min incubation for cecum extraction). Tissue was washed twice with cold complete media (RPMI containing 10% FBS, 20mM Hepes), minced with scissors, and then pieces were digested by 20-25 min incubation with complete media supplemented with 0.1 mg/ml Liberase (Roche 05401020001) and 0.1 mg/ml DNase 1 (Sigma, 10104159001) at 37°C incubator with shaking at 250 rpm. The digested cell suspension was homogenized using a 23G syringe, washed with complete media, and passed through a 70µ cell strainer. The pellet was resuspanded with 40% percoll (Percoll, Sigma P1644), and then 80% percoll was added gently underlay the 40% percoll. The samples in the gradient percoll were centrifuged at 860 g for 25 min at RT without acceleration and breaking. Interphase was collected for the immune enriched fraction. The cells were washed twice with DPBS. For bulk RNA extraction, see ‘RNA extraction’ in the method section.

#### Isolation of immune cells from mice spleen

Spleens from 5 naïve mice and 5 mice infected for 72 hours (with Salmonella SL1344) were collected for bulk RNA. Spleens were dissected and squashed against a 70 μm cell strainer into a tube with 5 ml of DPBS per spleen. The samples were then centrifuged (500g, 5 min in 4°c), washed twice with 5 ml cold DPBS, followed by immune cell enrichment by Lymphoprep (Lymphoprep, Alere Technologies AS). Lymphoprep step was adopted from Hen-Avivi and Avraham 2022^77^. For bulk RNA extraction, see ‘RNA extraction’ in the method section.

#### Isolation of immune cells from mice PP

PP from 5 naïve mice and 5 mice infected for 72h (with Salmonella SL1344) were collected for bulk RNA. PP from one naïve mouse and one 72h infected mouse were collected for single-cell RNAseq. PP from 5 naïve mice, 5 mice infected for 72h with Nal-WT Salmonella (IR715) and 4 mice infected for 72h with triple mutant Salmonella (FR90) were collected for bulk RNAseq and CFU. PP (one PP per sample, isolated from the 5 cm distal part of ileal tissue for RNAseq or middle part of the ileal tissue for CFU) were dissected and squashed against 70 μm cell strainer into a tube with 600 ul of DPBS for PP. The samples were then centrifuged (500g, 5 min in 4°c) washed with 600 ul cold DPBS and resuspended with cold DPBS. For bulk RNA extraction, see ‘RNA extraction’ in the method section. For single-cell RNA-seq of PP, see ‘Single-cell RNA libraries preparation’ in the method section. For CFU, see ‘Colony-forming units (CFU)’ in the method section.

### RNA preparation

#### RNA preparation from PBMCs samples

At the indicated time points after incubation, the samples were centrifuged (500g, 22°c, 5 min), the media was discarded, and the samples were washed with DPBS -/-. The samples were then resuspended with 5 mM EDTA in DPBS -/- and incubated for 5 min in RT, followed by gentle pipetting. This was done to detach cells that might attach to the dish. The samples were then centrifuged (500g, 22°c, 5 min) and resuspended in 350ul of RLT buffer (RNeasy Mini Kit, QIAGEN 74104) supplemented with 10ul beta-mercaptoethanol per 1ml RLT. Samples were kept at -80°c until RNA was extracted.

#### RNA preparation from mice tissue extracted cells

mice cells were resuspended in 350ul (ileum and cecum extracted cells) or 700ul (PP and spleen extracted cells) of RLT buffer (RNeasy Mini Kit, QIAGEN 74104) supplemented with 10ul beta-mercaptoethanol per 1ml RLT. Samples were kept at -80°c until RNA was extracted.

RNA for all samples (from PBMCs or mice tissue extracted cells) was extracted by RNeasy Mini Kit, QIAGEN 74104, including Dnase digestion (QIAGEN 79254). RNA samples concentration was measured by iQuant™ RNA HS Assay Kit (ABP Biosciences, N017), and RNA quality was measured by tapestation (High Sensitivity RNA ScreenTape, Agilent 5067-5579 and High Sensitivity RNA ScreenTape Sample Buffer, Agilent 5067-5580). RNA samples were kept at - 80°c until further use.

### Bulk RNA library preparation

RNA-seq libraries were prepared according to Cel-seq libraries protocol^82^ with a minor modification: the starting material was purified bulk RNA (up to 1ng RNA per library).

PBMCs under normoxia/ hypoxia were run on Illumina Miniseq instrument with MiniSeq Mid Output Kit (300 cycles) kit (FC-420-1004 Illumina) with a total coverage of ∼32 M reads and a median of ∼2.7 M reads per library. PBMCs naïve/ after *S.* Typhi infection were run on Illumina Miniseq instrument with MiniSeq Mid Output Kit (300 cycles) kit (FC-420-1004 Illumina) with a total coverage of ∼8.3 M reads and a median of ∼350,000 reads per library. Mice samples from naïve and infected ileum were run on Illumina NovSeqX instrument with NovaSeq 6000 S1 Reagent Kit v1.5 (100 cycles) kit (20028319, Illumina) with a total coverage of ∼220 M reads and a mean of ∼9 M reads per library. Mice samples from naïve and infected cecum were run on Illumina NovaSeq 6000 instrument with NovaSeq 6000 SP Reagent Kit v1.5 (500 cycles) kit (20028402, Illumina) with a total coverage of ∼82 M reads and a mean of ∼6.5 M reads per library. Spleen and PP from naïve and infected ileum were run on Illumina NovaSeq 6000 instrument with NovaSeq 6000 SP Reagent Kit v1.5 (500 cycles) kit (20028402, Illumina) with a total coverage of ∼192 M reads and a mean of ∼4.8M reads per library. PP from naïve, IR715-WT infected and triple mutant (FR90) infected mice were run on Illumina NovSeqX instrument with NovaSeq 6000 S1 Reagent Kit v1.5 (100 cycles) kit (20028319, Illumina) with a total coverage of ∼20 M reads and a mean of ∼1 M reads per library.

### Single-cell RNA libraries preparation

#### For 18 samples from the human typhoid infection model

PBMCs from 6 individuals (3-TD, 3-nTD) from 3-time points per individual were defrosted, counted with viability staining (ViaStain™ AOPI Staining Solution, Revvity CS2-0106-5ml), resuspended in RPMI+ media (RPMI 1640 with L-Glutamine supplemented with 10% heat inactivated fetal bovine serum and 1 mM sodium pyruvate), plated on U-shape 96-well-untreated plates (5* 10^5 cells/well, one well per individual) and incubated at 37°c and 5% CO_2_ over-night. After overnight incubation, samples were centrifuged (500g, 22°c, 5 min), the media was discarded, and the samples were washed with DPBS -/-. Detaching the cells from the dish was done by resuspension with 5 mM EDTA in DPBS -/- followed by 5 min incubation in RT. After the incubation, the cells were gently pipetted. The samples were then centrifuged (500g, 22°c, 5 minutes) and resuspended in DPBS. Each sample was counted with viability staining (ViaStain™ AOPI Staining Solution, Revvity CS2-0106-5ml) and resuspended with 0.04% BSA (UltraPure™ BSA (50 mg/mL), AM2616 Thermo Fisher Scientific) in DPBS, and directly used for single-cell sequencing by the Chromium Single Cell 3′ Reagent version 3 kit and Chromium Controller (10X Genomics, CA, USA). Each sample from the 18-PBMCs samples was run in a separate 10X lane. Library quality and concentration were assessed according to the manufacturer’s instructions. The libraries were run with 6 Nextseq kits (NextSeq® 500 High Output v2 Kit (75 cycles), Illumina FC-404-2005) on Nextseq instrument (Illumina), with a total coverage of ∼3700 M reads.

#### For mouse PP samples

cells extracted from mice PP (one naïve mouse, one infected mouse) were counted with viability staining (ViaStain™ AOPI Staining Solution, Revvity CS2-0106-5ml) and resuspended with 0.04% BSA (UltraPure™ BSA (50 mg/mL), AM2616 Thermo Fisher Scientific) in DPBS, barcoded with hashing antibodies (TotalSeq B hashing indexes (BioLegend, 199902), hash index TGCCTATGAAACAAG for naïve PP and hash index CCGATTGTAACAGAC for infected PP at 72 hpi) according to the manufacturer’s instructions. The samples were then directly used for single-cell sequencing by the Chromium Single Cell 3′ Reagent version 3 kit and Chromium Controller (10X Genomics, CA, USA), and run on one 10X lane. Library quality and concentration were assessed according to the manufacturer’s instructions. The libraries were run with with NovaSeq 6000 S1 Reagent Kit v1.5 (100 cycles) kit (20028319, Illumina) on NovaSeqX instrument (Illumina) with a total coverage of ∼1246 M reads

### Colony-forming units (CFU)

PP isolated from 5 cm middle part of ileal tissue (PP from the 5 cm distal part of these mice were used for bulk RNAseq) from 5 naïve mice, 5 Nal-WT Salmonella (IR715)-infected mice for 72h, and 4 triple mutant Salmonella (FR90) infected mice for 72h were collected and processed as appears in ‘Mice tissue extraction’ in the method section. The cells were then suspended with 0.1% triton X-100, pipetted vigorously to lyse them, and incubated at room temperature for 10 min. Serial dilutions were made in DPBS, plated on LB + 50 µg/ml nalidixic acid plates, grown overnight in 37°c, and counted with Scan® 500 automatic colony counter. Reagents used for the experiment (LB, DPBS, and 0.1% triton X-100) were also plated on LB agar plates for controls.

## Bioinformatics analysis

### scRNA-seq of PBMCs samples from the S. Typhi human challenge model

#### Processing

The Cell Ranger Single-Cell Software Suite (https://support.10xgenomics.com/single-cell-gene-expression/software/pipelines/latest/what-is-cell-ranger) was used to perform alignment to the genome (GRCh38), barcode assignment for each cell, and gene counting by unique molecular identifier (UMI) counts for each sample. Each sample was run on a different lane (total of 18 lanes), and samples from all individuals and time points were merged together using cell ranger aggr, which aggregates multiple GEM wells. Overall, we sequenced 165,415 cells with a median of 9267 cells per sample, ∼8800 mean reads per cell, ∼1600 median UMI count per cell, and ∼600 median genes per cell.

#### QC filtration and normalization

Only genes with at least one UMI count detected in at least one cell were used. Cells with a high percentage of mitochondrial genes (more than 10%) were excluded. All cells had at least a total count of 200 UMIs. Data was normalized to a library size factor, with factors calculated by dividing the total UMI counts in each cell by the median of the total UMI counts across all cells. Normalized data was log2-transformed. Cell cycle and ribosomal genes were filtered out prior to the selection of variable genes. Variable genes were selected based on fitting the data to a simple noise model based on the gene’s mean expression and dispersion (coefficient of variance) across all single cells.

#### Clustering and data analysis

Principal component analysis (PCA) was performed on the variable genes, and the first 20 PCs were selected for downstream analysis based on the inflection point of the curve of the variance explained by each successive principal component. t-distributed stochastic neighbor embedding (tSNE) in the PC space was calculated for the visualization of the data. The Euclidian distance in the PC space and the Jaccard similarity were used to build a k-nearest neighbor (KNN)-graph with k=20. Unsupervised clustering of the single cells was done using Louvain community detection. We used an iterative implementation of the Louvain method^83^ to optimize modularity-like quality function by iterating the Louvain algorithm until convergence. Cluster identity was inferred using cluster-specific genes and cell type marker genes, and all clusters from the same cell type were merged to get the cell type level clustering with NK cells, T cells, B cells, monocytes and macrophages, and DCs (**Supplementary Fig. 3a**). Next, we re-cluster each cell type to the subset level using Louvain algorithm, and thereafter merged subsets which similar expression patterns. Overall, we obtained 9 NK subsets, 15 T cells subsets, 13 B cells subsets, 15 monocyte and macrophage subsets, and one subset of DCs (**Supplementary Fig. 4a and Supplementary Fig. 5a**). Subsets classification and annotation were based on subsets marker genes and immune and metabolic subset-specific genes.

### Local inverse Simpson’s Index (LISI) score

The LISI score^84^ was used to assess the mixing of single cells across individuals and disease outcomes. The score is calculated based on the local neighbors on the tSNE embedding. We used the integrated LISI (iLISI) implementation^34^, with scores ranging from 1 (no mixing) to the maximal number of categories in the variable of interest, representing perfect mixing. This corresponds to a score of 6 for mixing across individuals and 2 for mixing across disease outcomes. By comparing the mixing scores across individuals to those across disease outcomes for each cell, we quantified the extent to which variability in immune cell responses is driven by infection outcomes relative to inter-individual variability.

### Pathway enrichment analysis and Gene Set Enrichment Analysis (GSEA)

Enrichment analysis of significant DEGs in KEGG or MSigDB pathways was performed using a hypergeometric test with FDR correction for multiple tests. GSEA was conducted by ranking genes based on their fold-change between the two tested conditions and calculating the enrichment scores (ES), maximum ES, and normalized ES (NES)^85^. P-values were determined from the null distribution of maximum ES values obtained through 1,000 random permutations. The group of enriched genes for each significant pathway was defined as those with a fold-change greater than the fold-change at the position of the maximum ES.

### Monocytes infection-induced modules from the scRNA-seq data

We identified all genes that were significantly upregulated in the monocytes and macrophages following *S.* Typhi challenge, applying a 1% FDR and 1.1fold-change cutoff for comparisons between T0 versus T12h or T0 versus T7d/TD. These genes were then clustered across all monocyte and macrophage subsets to identify co-regulated gene modules (**Supplementary Fig. 4b**). The co-regulated modules were categorized into immune-related pathways based on the functions of the genes in each module and their enrichment to KEGG and MSigDB pathways (**Supplementary Table 5**).

To analyze the dynamics of infection-induced modules across time points in nTD and TD individuals, we calculated a per-cell score for each module. These scores were aggregated for each individual, and then mean and standard error (SE) were calculated for nTD and TD groups. The per-cell score for each module was defined as the mean z-score of all genes in the module, normalized to a range of 0 to 1 (by subtracting the minimum value across all cells and dividing by the range between the maximum and minimum values). To compute a p-value for the difference in dynamics between nTD and TD groups, we calculated the difference between the integrals of the nTD and TD curves for each module and generated a random model using 1,000 permutations.

### Monocytes metabolic modules from the scRNA-seq data

We extracted all metabolic pathways from the KEGG database^86^, comprising 85 metabolic pathways with 1696 genes, and added a hypoxia pathway from Tawk et al., 2016^87^, which previously analyzed several hypoxia signatures from different cancer patients. To focus specifically on the hypoxia pathway independent of glycolysis, we removed glycolysis-associated genes from the hypoxia pathway gene list, resulting in 51 genes (**Supplementary Table 10**). The metabolic genes were then clustered across all monocyte and macrophage subsets. For each co-regulated group of metabolic genes, we calculated enrichment scores for all metabolic pathways. A co-regulated group of genes significantly enriched for a specific metabolic pathway was defined as a metabolic module (**Supplementary Fig. 5d**).

### Sliding window analysis

To assess the relationship between the expression of two gene-list signatures, we calculate an average trend line using a sliding window approach. The x-axis was divided into bins of size 0.01, and for each bin, the mean expression of the y-axis signature was calculated based on the corresponding y-axis values within this bin. These calculated means were then used to generate the average line.

For the analysis of glycolysis versus OXPHOS, we observed two distinct behaviors at low and high glycolysis rates: a positive correlation at low rates and a negative correlation at high rates. To separate between these two behaviors, we computed the derivate of the average line at each position. The transition point between low and high glycolysis rates was defined as the position where the derivate equaled zero.

### Hypoxic subsets six-genes signature

To generate a signature that captures all hypoxic subsets independent of cell type, we analyzed the expression of marker genes from each hypoxic subset across all subsets from all cell types. We selected genes that were expressed in all four hypoxic subsets (NK, T, B, and monocyte subsets) but exhibited low expression levels in all other subsets.

### Bulk RNA-seq processing and normalization

The cel-seq pipeline (https://github.com/yanailab/CEL-Seq-pipeline) was used for sample demultiplexing, alignment to the genome (GRCh38 or mm10), and gene counting for each experiment independently to generate an expression matrix. The expression matrix was normalized to a library size factor by dividing the total number of reads from each sample by the median of the total number of reads across all samples in the experiment. Normalized data was log2-transformed.

### Bulk RNA-seq of PBMCs incubated in a hypoxic chamber or ex vivo infected with S. Typhi

One sample was excluded from the analysis due to low coverage (*ex vivo* infected PBMCs at 24 hpi replicate 1), a minimal expression threshold of 3 was set for the normalized expression matrix. DEGs following hypoxia (15 or 22h) or *ex vivo* infection (4, 8, or 24 hpi) were calculated using a two-samples t-test compared to the matched naïve samples for each condition and FDR correction. The HIF1α targets were curated by combination of several HIF1α signatures from MSigDB and KEGG (see **Supplementary Table 10**). The heat map in **Supplementary Fig. S6a** showing the expression levels of all HIF1α targets curated list that were significantly differentially expressed due to hypoxia or following *ex vivo* infection with 20%FDR.

### A cohort of sepsis patients

We analyzed the expression of our six-gene hypoxia signature in a public cohort of sepsis patients^46^ (GSE46955). This cohort contains bulk RNA expression of blood monocytes isolated from gram-negative sepsis patients during sepsis and following their recovery and from healthy donors as control. We downloaded the log2-transformed normalized data and analyzed the expression of our six-gene hypoxia signature. We used only the unstimulated samples from this cohort.

### A cohort of Crohn’s disease patients

We analyzed the expression of our six-gene hypoxia signature in a cohort of Crohn’s disease patients^47^. The cohort contains scRNA-seq data from the terminal ileal biopsies of 50 Crohn’s disease patients with different disease severity and 71 healthy controls. The data is not publicly available but can be viewed using the IBD portal (https://www.ibd-cell-portal.org). We checked the expression of our six-genes hypoxia signature and identified that all of them are co-expressed from the same cells. We then defined a signature expression of our six genes using the portal and explored the expression of the signature across the different attributes available in the portal, such as the cell type, disease state, and disease severity.

### Bulk RNA-seq of human cutaneous leishmaniasis lesions

We tested the expression of genes significantly induced in hypoxic lesions of cutaneous leishmaniasis relative to non-hypoxic lesions in our human data from the *S.* Typhi challenge model. We extracted the gene list from Fowler et al., 2024^50^, which defined a list of 44 genes that were significantly expressed higher in hypoxic lesions relative to non-hypoxic lesions with 1%FDR and above 1.5fold difference.

### Mouse typhoid model bulk RNA-seq of ileum, cecum, spleen, and PP

#### QC filtration

Four samples out of 44 were excluded from the analysis due to low coverage: 1) a sample from the ileum at 72 hours post-infection (hpi), 2) a naïve sample from the cecum, 3) a sample from the cecum at 24 hpi, and 4) a sample from the PP at 72 hpi. The data was normalized for each tissue alone, only for the analysis of all tissues together all data was normalized together. A minimal expression threshold was set to 2 for each normalized expression matrix.

#### Analysis

Differentially expressed genes (DEGs) between tissues were calculated using one-way ANOVA and false discovery rate (FDR) correction. DEGs before and after infection for each tissue alone (naive versus 24 hpi and naïve versus 72 hpi) were calculated by a two-sample t-test and FDR correction.

### scRNA-seq data from the PP of naïve and infected mice in the typhoid model

#### Processing

The Cell Ranger Single-Cell Software Suite (https://support.10xgenomics.com/single-cell-gene-expression/software/pipelines/latest/what-is-cell-ranger) was used to perform alignment to the genome (mm10), barcode assignment for each cell, gene counting by unique molecular identifier (UMI) counts, and samples demultiplexing based on the TotalSeq B hashing indexes. Overall, we sequenced 2525 cells, 661 cells from the naïve mouse and 1864 cells from the infected mouse, with ∼70,000 mean reads per cell, ∼2000 median UMI count per cell, and ∼5500 median genes per cell.

#### QC filtration and normalization

Only genes with at least one UMI count detected in at least one cell were used. Cells with a high percentage of mitochondrial genes (above 10%) were excluded. All cells had at least a total count of 500 UMIs. Data was normalized to a library size factor, with factors calculated by dividing the total UMI counts in each cell by the median of the total UMI counts across all cells. Normalized data was log2-transformed. Variable genes were selected based on fitting the data to a simple noise model based on the gene’s mean expression and dispersion (coefficient of variance) across all single cells.

#### Data analysis

Principal component analysis (PCA) was performed on the variable genes, and the first 54 PCs were selected for downstream analysis based on the inflection point of the curve of the variance explained by each successive principal component. tSNE in the PC space was calculated for data visualization. The Euclidian distance in the PC space and the Jaccard similarity were used to build a KNN graph with k=30. Unsupervised clustering of the single cells to the cell type level was done using Louvain community detection on the KNN-graph. Cluster identity was inferred using cluster-specific genes and known cell-type marker genes.

### Bulk RNA-seq of naïve and S. Typhimurium wild-type-infected and triple mutant-infected PP

#### QC filtration

Two samples out of 14 samples were excluded from the analysis due to a high percentage of mitochondrial genes (above 18%; one naïve sample and one mutant-infected sample). An additional sample of wild-type-infected PP was excluded from the analysis due to high expression of epithelial genes relative to other replicates (probably was not enriched as expected for immune cells). A minimal expression threshold for the normalized data was set to 3.

### Bulk RNA-seq of naïve and Yersinia pseudotuberculosis-infected PP

We analyzed a public dataset of bulk RNA-seq of mice infected with *Yersinia pseudotuberculosis*^18^. The dataset contains PP extracted from three naïve mice and three infected mice for 72 hours. We downloaded the raw data and normalized it to a library size factor by dividing the total number of reads from each sample by the median of the total number of reads across all samples. Normalized data was log2-transformed, and a minimal expression threshold was set to 3. PCA and GSEA were performed to identify the significantly expressed pathways between naïve and infected PP in mice.

## Notes

### Competing Interest Statement

The authors have declared no competing interest.

