## Supplementary Figures 1-7 for "Hypoxia-related immune subsets induced by *Salmonella* Typhi infection link early bacterial gut invasion to human infection outcomes"

### Supplementary Figure 1

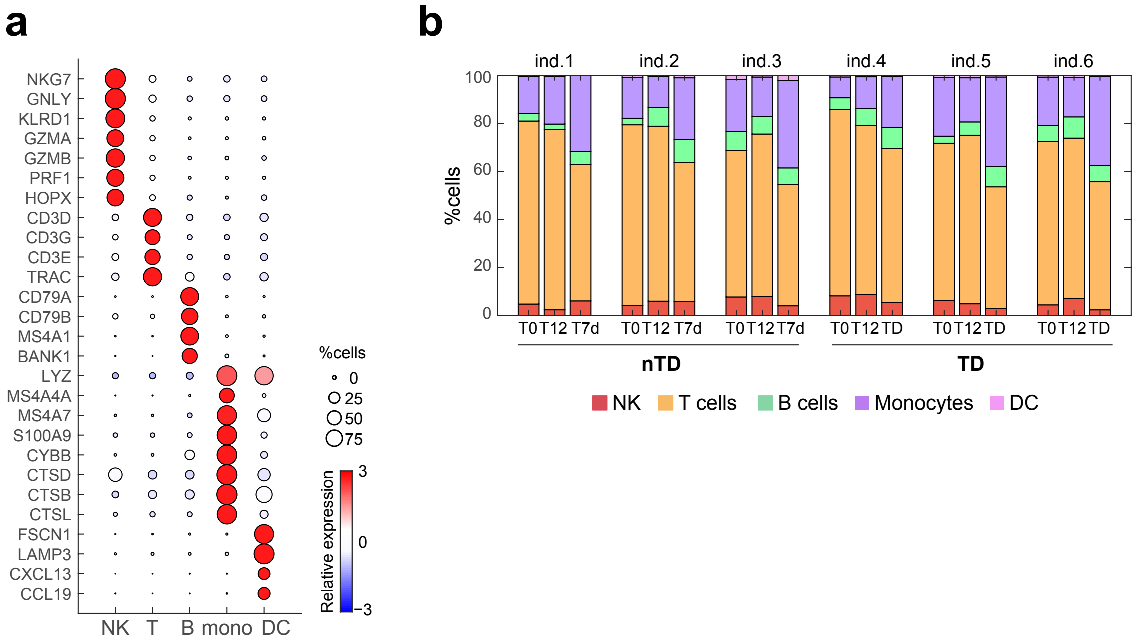


**Supplementary Figure 1: Dynamics of PBMC cell-type composition in the *S.* Typhi human challenge model. (a)** Dot plot showing marker gene expression levels for each cell type. Dot size represents the percentage of cells expressing a given gene within each cluster, and dot color indicates relative expression levels (see legend on the right). **(b)** Cell-type composition of PBMC samples for each individual during *S.* Typhi challenge. Color coding corresponds to cell types listed in the legend at the bottom.

### Supplementary Figure 2

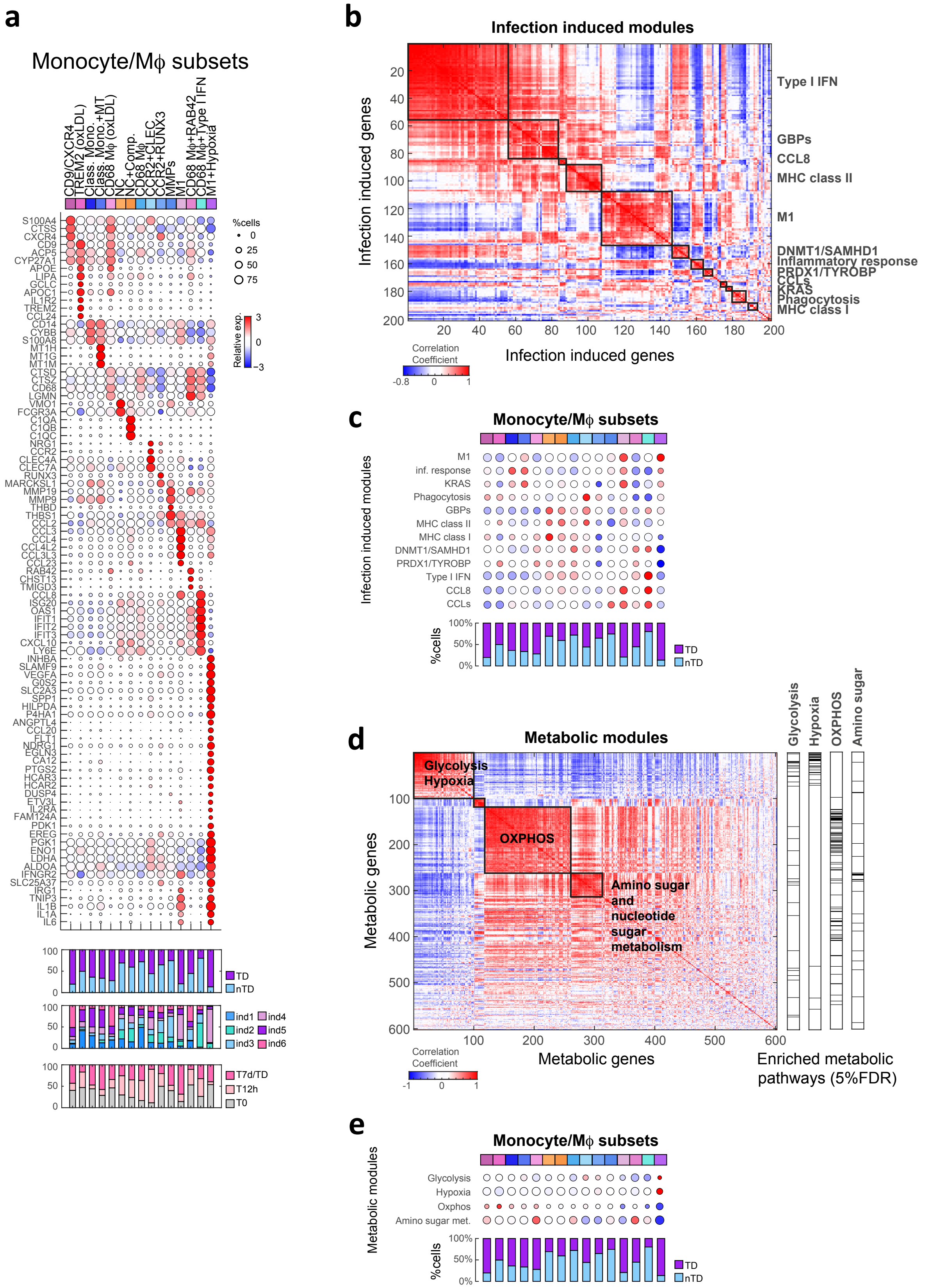

**Supplementary Figure 2: Infection-induced and metabolic states of monocyte and macrophage subsets in the *S.* Typhi human challenge model. (a)** Dot plot showing expression levels of marker genes across monocyte and macrophage subsets. Dot size represents the percentage of cells in a subset expressing a given gene, and dot color indicates relative expression levels (see legend on the right). Bar plots below the dot plot display the composition of disease outcomes (nTD and TD), individuals (ind1-ind6), and time points (T0, T12h, and T7d/TD) within each subset. **(b)** Correlation matrix of infection-induced genes across monocyte and macrophage subsets. Black squares indicate co-regulated groups of infection-induced genes, referred to as infection-induced modules. Annotations for each module are shown on the right. **(c)** Dot plot showing the expression levels of infection-induced modules across all monocyte and macrophage subsets. Dot size represents the percentage of cells in a subset expressing a given module, and dot color indicates relative expression levels (legend as in (a)). The bar plot below represents the disease outcome composition for each subset. **(d)** Correlation matrix of metabolic genes across monocyte and macrophage subsets. Black squares indicate co-regulated groups of metabolic genes, which are defined as metabolic modules if enriched (%5FDR) for a specific metabolic pathway. Bars on the left highlight the location of the metabolic genes associated with each enriched pathway. **(e)** Dot plot showing the expression levels of metabolic modules across all monocyte and macrophage subsets. Dot size represents the percentage of cells in a subset expressing a given module, and dot color indicates relative expression levels (legend as in (a)). The bar plot below represents the disease outcome composition for each subset.

### Supplementary Figure 3

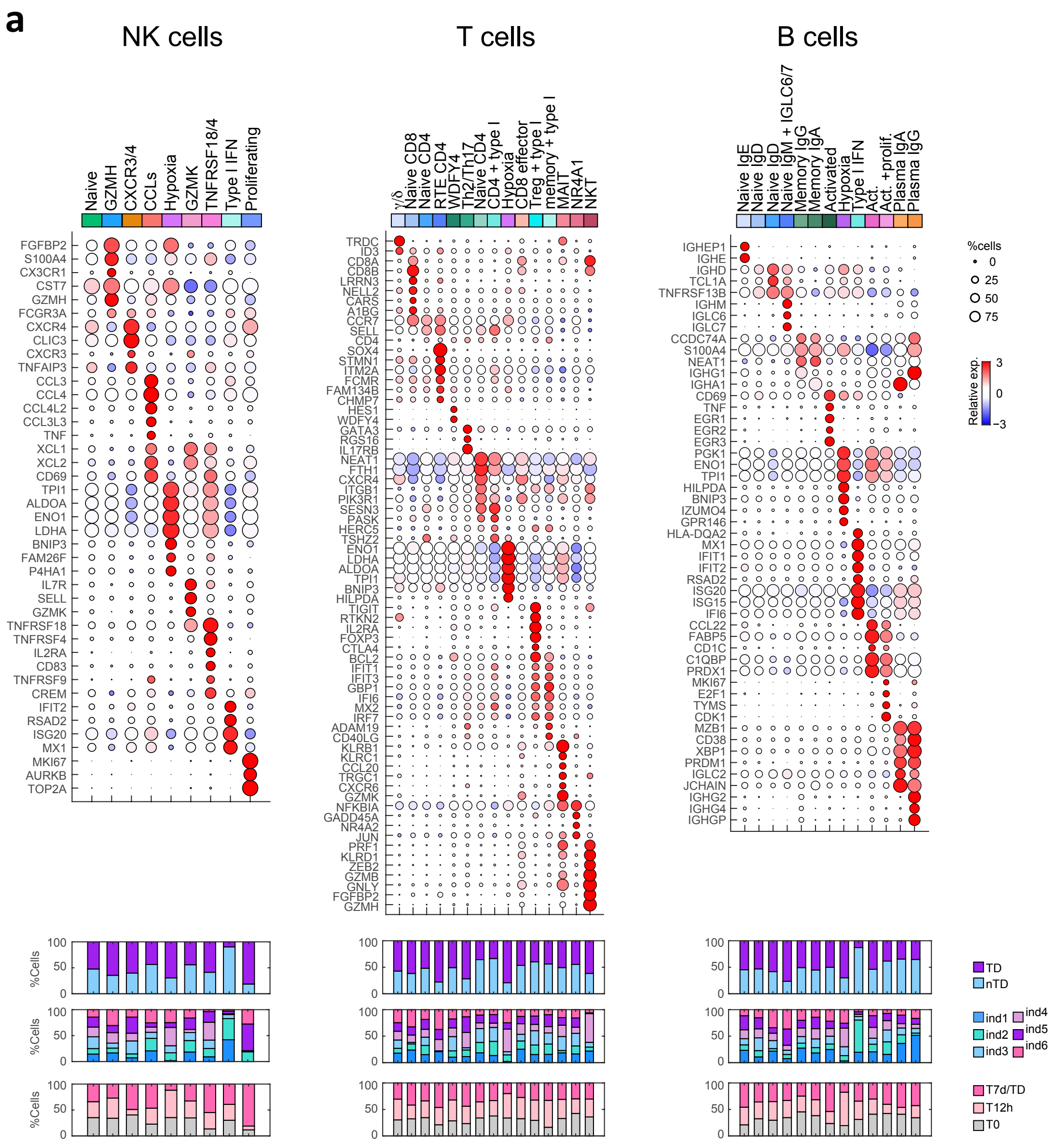


**Supplementary Figure 3: Repertoire of circulating immune cells in individuals from the S. Typhi human challenge. (a)** Dot plot showing expression levels of marker genes across NK subsets (left), T cell subsets (middle), and B cell subsets (right). Dot size represents the percentage of cells in a subset expressing a given gene, and dot color indicates relative expression levels (see legend on the right). Bar plots below the dot plots display the composition of disease outcomes (nTD and TD), individuals (ind. 1-6), and time points (T0, T12h, and T7d/TD) within each subset.

### Supplementary Figure 4

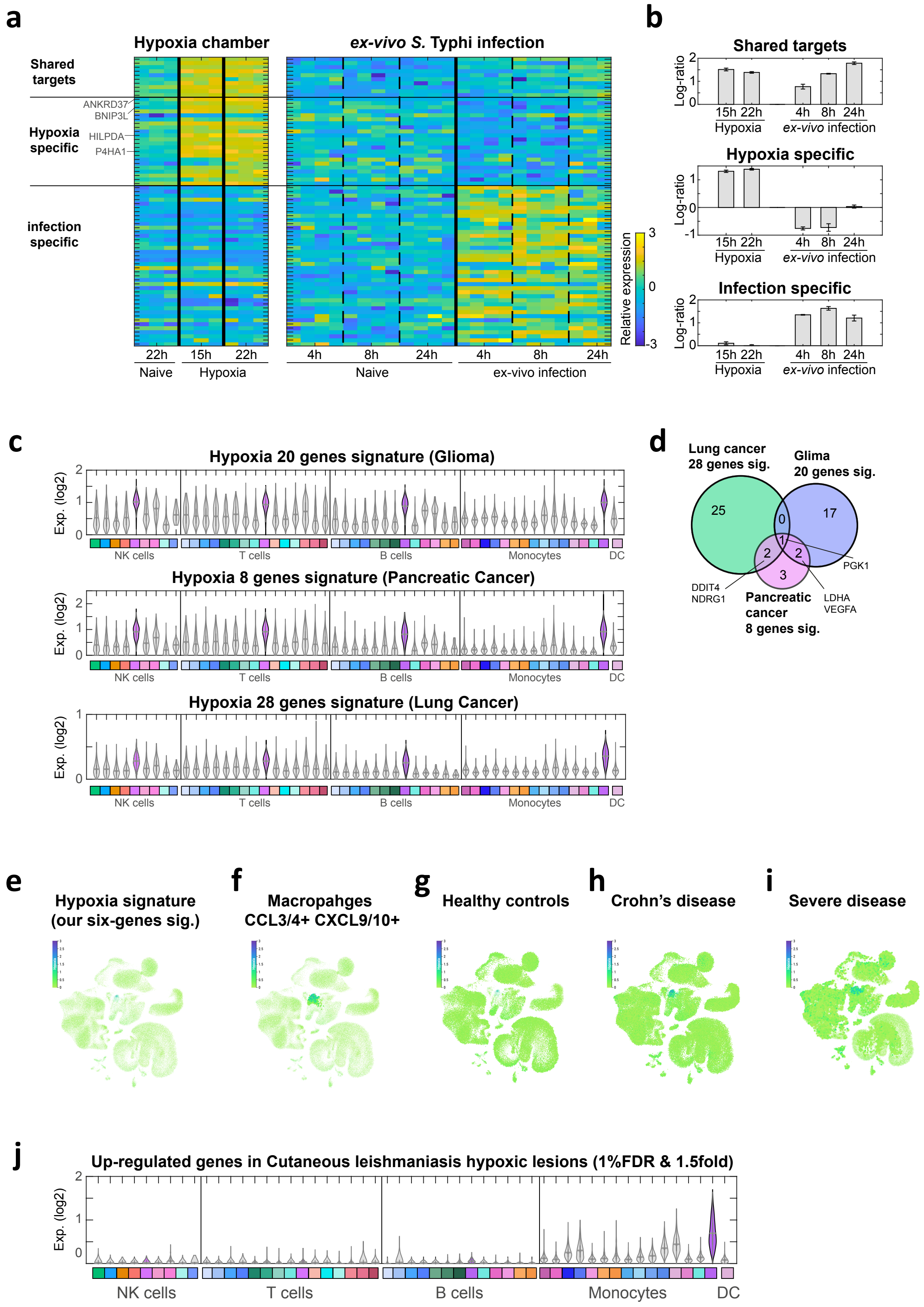

**Supplementary Figure 4: The hypoxia gene signature and hypoxic subsets are associated with infectious disease-related pathologies. (a)** Heat map showing expression levels of HIF1α target genes under hypoxic conditions (15 or 22 hours (h) in a hypoxia chamber), or following *ex vivo* infection with *S.* Typhi for 4h, 24h, or 48h. The heat map is divided into three categories: 1) Shared targets induced under both hypoxia and infection, 2) Hypoxia-specific targets induced only under hypoxic conditions, and 3) infection-specific targets induced only following *S.* Typhi infection. **(b)** Bar plots representing the fold change of genes within each category (shared, hypoxia-specific, infection-specific) under hypoxia or *ex vivo* infection conditions, relative to matched naïve conditions. Mean and standard error (SE) values are presented. **(c)** Violin plots showing the expression levels of hypoxia signatures identified in cancer studies (glioma (Lin et al., 2020), pancreatic cancer (Abou Khouzam et al., 2021), and lung cancer (Lane et al., 2022)) across immune subsets from the *S.* Typhi human challenge model. The mean expression of each signature is shown for all subsets, with hypoxic subsets highlighted in purple. **(d)** Venn diagram showing the overlap among the three hypoxia signatures from (c). **(e-i)** Analysis of the six-gene hypoxia signature (from Fig. 4a) in a cohort of Crohn's disease patients (Krzak et al., 2023; data obtained from the IBD portal https://www.ibd-cell-portal.org). Uniform Manifold Approximation and Projection (UMAP) of immune, stromal, and epithelial cells from terminal ileum biopsies of Crohn's disease patients is shown. Expression levels of the six-gene hypoxia signature are depicted with a color bar for relative expression. Highlighted cells represent specific attributes: **(e)** None of the cells were selected, cells expressing the signature are limited to a specific subset. **(f)** Macrophages expressing CCL3/4 and CXCL9/10 are selected and highlighted, cells from this population show high expression levels of the six-gene hypoxia signature. **(g)** Cells from healthy controls are highlighted, cells from this attribute do not express the six-gene hypoxia signature. **(h)** Cells from Crohn's disease patients are highlighted, cells from this attribute show high expression of the six-gene hypoxia signature. **(i)** Cells from Crohn's disease patients with severe disease are highlighted, cells from this attribute show high expression of the six-gene hypoxia signature. **(j)** Violin plot showing the expression levels of genes up-regulated in cutaneous leishmaniasis hypoxic lesions compared to non-hypoxic lesions (1%FDR and 1.5fold-change; Fowler et al., 2024), across immune subsets from the *S.* Typhi human challenge model. Presented violin plot with the mean expression of the genes in each cell across all subsets; hypoxic subsets are highlighted in purple.

### Supplementary Figure 5

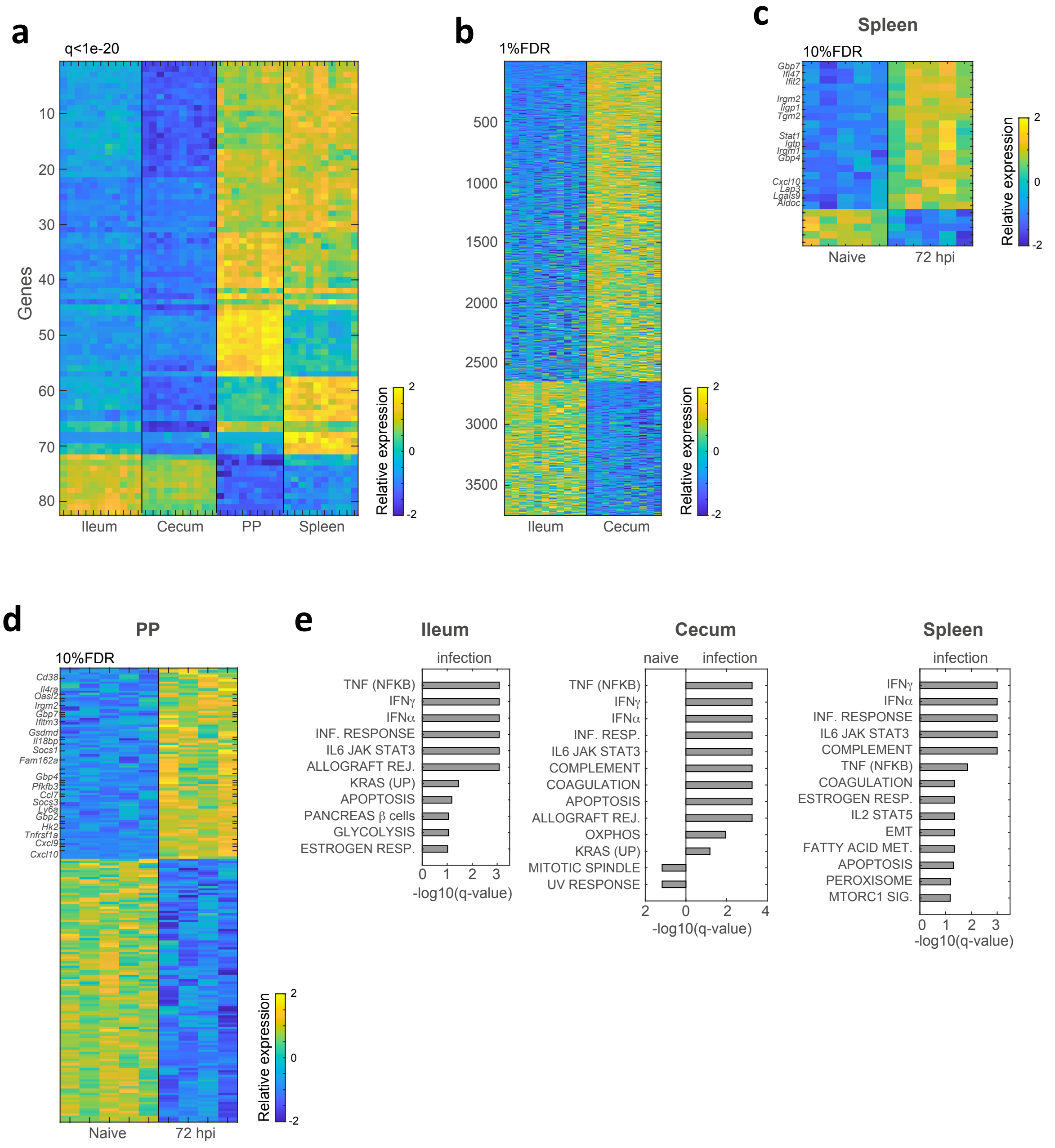


**Supplementary Figure 5: Transcriptional changes of immune cells in the gut in a mouse model of typhoid fever. (a)** Heatmap showing the expression levels of the significant DEGs between the different tissues: ileum, cecum, PP, and spleen, independent of time-point (one-way ANOVA, FDR<$1x10^{-20}$). Samples from all time points are included for each tissue. **(b)** Heatmap showing the expression levels of significant DEGs between ileum and cecum, independent of time-point (two samples t-test, 1%FDR). Samples from all time points are presented. **(c)** Heatmap showing expression levels of significant DEGs between naïve and infected spleens at 72 hpi (two-samples t-test, 10%FDR). **(d)** Heatmap showing expression levels of significant DEGs between naïve and infected PP at 72 hpi (two-sample t-test, 10%FDR). **(e)** GSEA for *S.* Typhimurium infected ileum, cecum or spleen at 72 hpi compared to naïve tissue.

### Supplementary Figure 6

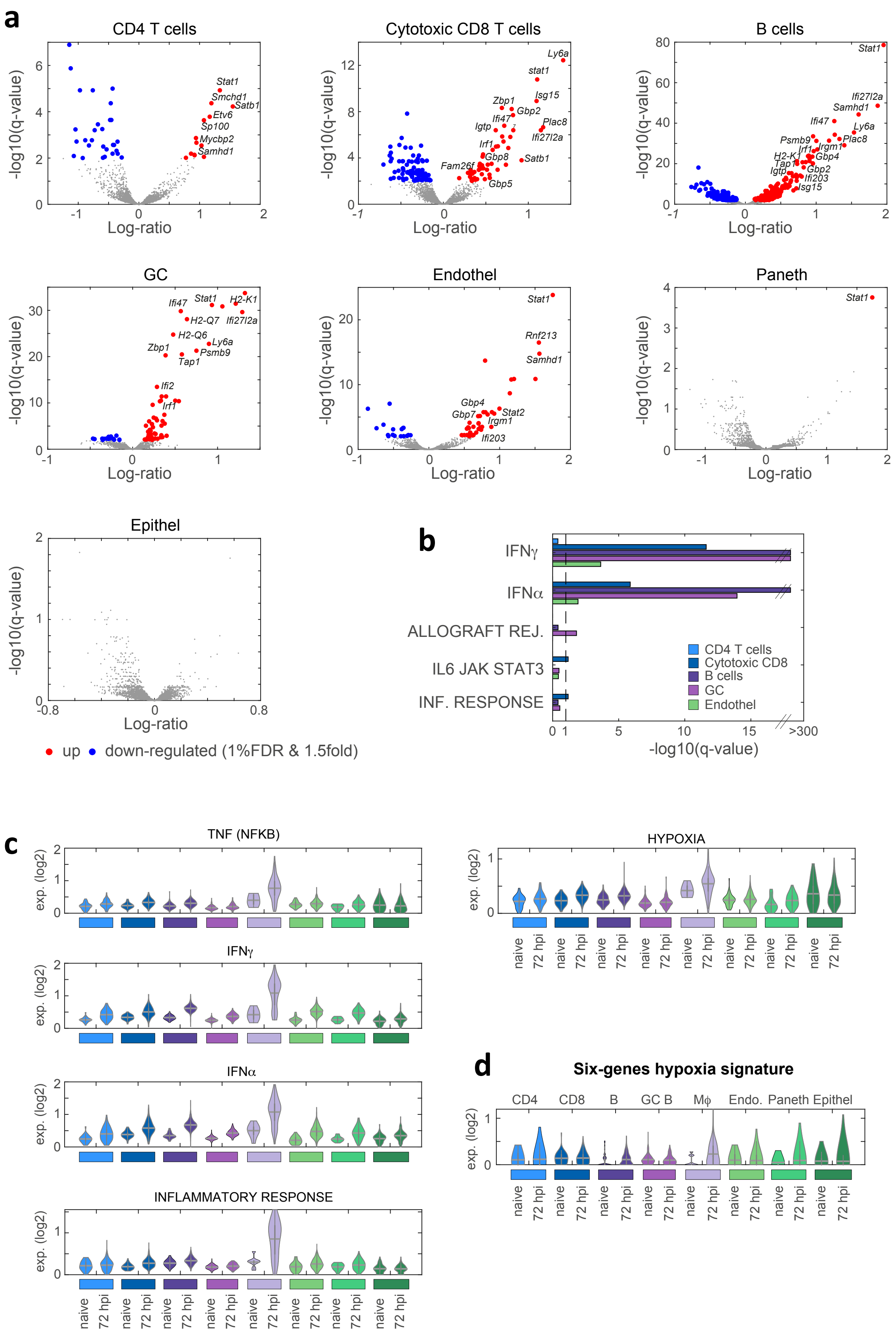

**Supplementary Figure 6: Differentially expressed genes and pathways in the PP of infected versus naïve mice in the typhoid fever model. (a)** Volcano plots displaying gene expression differences between naïve and infected PP at 72 hpi for each cell type. Red and blue circles indicate significantly upregulated and downregulated genes in infected PP, respectively (1%FDR and 1.5-fold change, two-sample *t-test*. **(b)** Pathway enrichment for the significant DEGs in each cell type. Color coding for cell types is shown in the legend. **(c)** Violin plots displaying the expression distribution of genes from the top significant immune and metabolic pathways induced in infected PP at 72 hpi (from Fig. 5g). Expression is shown across clusters (cell types), separated for naïve and infected cells. (**d)** Violin plot showing expression levels of the six-gene hypoxia signature across immune cell types from naïve and infected PP (as is (c)).

### Supplementary Figure 7

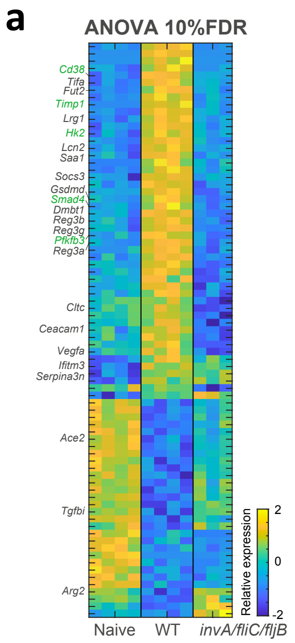


**Supplementary Figure 7: The hypoxia gene signature in the PP of infected mice depends on PP invasion and colonization. (b)** Heat map depicting genes significantly differentially expressed (10%FDR, one-way ANOVA) among naïve, wild-type infected, and *invA/fliC/fljB* mutant infected PP at 72 hours post-infection. Selected gene names are shown, with hypoxia-related genes highlighted in green.
